# Resolving inter-regional communication capacity in the human connectome

**DOI:** 10.1101/2022.09.28.509962

**Authors:** Filip Milisav, Vincent Bazinet, Yasser Iturria-Medina, Bratislav Misic

**Affiliations:** McConnell Brain Imaging Centre, Montréal Neurological Institute, McGill University, Montréal, QC, Canada

## Abstract

Applications of graph theory to the connectome have inspired several models of how neural signaling unfolds atop its structure. Analytic measures derived from these communication models have mainly been used to extract global characteristics of brain networks, obscuring potentially informative interregional relationships. Here we develop a simple standardization method to investigate polysynaptic communication pathways between pairs of cortical regions. This procedure allows us to determine which pairs of nodes are topologically closer and which are further than expected on the basis of their degree. We find that communication pathways delineate canonical functional systems. Relating nodal communication capacity to meta-analytic probabilistic patterns of functional specialization, we also show that areas that are most closely integrated within the network are associated with higher-order cognitive functions. We find that these regions’ proclivity towards functional integration could naturally arise from the brain’s anatomical configuration through evenly distributed connections among multiple specialized communities. Throughout, we consider two increasingly constrained null models to disentangle the effects of the network’s topology from those passively endowed by spatial embedding. Altogether, the present findings uncover relationships between polysynaptic communication pathways and the brain’s functional organization across multiple topological levels of analysis and demonstrate that network integration facilitates cognitive integration.

## INTRODUCTION

The anatomical connectivity of neural circuits supports signal propagation between neuronal populations [8]. Signals, in the form of electrical impulses, are relayed via axonal projections (monosynaptic communication). Wiring among multiple populations forms circuits in which signals can also be relayed between populations that do not share a direct projection, but can be reached via multiple synapses (polysynaptic communication) [81]. Thus, the architecture of the brain’s connectome shapes communication patterns and integration among specialized brain regions [7, 134].

The conventional approach to studying communication in brain networks is to model the global capacity of the network. Broadly, this paradigm involves estimating communication efficiency between all pairs of regional nodes and then taking the average to summarize communication efficiency with a scalar value [1, 44, 65, 130]. For instance, the oft-studied global efficiency statistic is defined as the inverse of the mean shortest path length among all pairs of nodes in a network [20, 75]. However, this broad, globally-focused approach obscures potentially informative heterogeneity of communication between specific pairs of regions. Yet, there is increasing appreciation for local heterogeneity in the brain, including spatial patterning of microarchitecture [23, 43, 57, 62, 136], dynamics [113, 137], and functional specialization [56, 142]. Importantly, numerous studies have reported evidence of regional heterogeneity in connection profiles or fingerprints of regions [83], as well as patterns of structure-function coupling [14, 79, 104, 133, 145] and electromagnetic-haemodynamic coupling [112].

Previous studies have also considered inter-regional communication capacity in specific settings. Most efforts have been focused on the predictive utility of pairwise communication measures, relating them to functional connectivity or behavior [45, 49, 66, 110, 133, 145]. Other studies have investigated its potential in distinguishing between patients with neurological disorders and healthy controls [30, 31, 76, 77]. The most widely studied communication mechanism is the topological shortest path [7], but numerous other models have been proposed, involving both routing protocols relaying signals through specific paths and diffusive processes in which neural signaling is driven by local network features [6, 30, 45, 89–91, 111]. For example, communicability, a communication measure integrating all possible walks on a network [30, 39], has been used to cluster patients [30, 31, 76, 77], characterize regional lesion effects [4, 31, 77], and even compensatory inter-regional responses to Alzheimer’s disease [76]. Nevertheless, global aggregated accounts of the brain’s communication capacity remain the norm and are often used even for diffusive processes, e.g., mean navigation time/efficiency [63, 64, 72, 111], diffusion efficiency [5, 25, 44, 46], and average communicability [114]. Furthermore, while it is commonplace to provide a null frame of reference when evaluating the prominence of global network attributes [132], this procedure is not applied at the inter-regional level. In summary, how inter-regional and regional communication preferences are organized remains poorly understood but methodologically accessible.

Here we develop a simple method to study the capacity for pairs of brain regions (dyads) to communicate with each other. We deconstruct the conventional global approach and estimate communication capacity without averaging over pairs of regions. In addition, we introduce a procedure to standardize the communication capacity between pairs of regions by their communication capacity in a population of rewired null networks, allowing us to identify pairs of regions with greater or less than expected communication capacity. An important advantage of this method is that it can be used in combination with any null network model. Increasingly constrained surrogates constitute increasingly conservative benchmarks which more closely resemble the empirical network under study [132]. Using them in parallel can simultaneously allow us to control for a covariate or disentangle its effect from those of other unconstrained features in a more liberal null model [132]. Degree-preserving null models are the most commonly used for network statistic normalization [61, 85, 92, 127, 131]. They allow to mitigate the effect of this simple, but influential graph feature to assess to which extent a network attribute is unexpected in contrast to a random graph which only preserves the empirical network’s degree sequence. In doing so, they highlight the effect of more subtle topological properties on a given statistic. Here, the main null model under consideration further constrains the empirical network’s weighted degree sequence using a simulated annealing procedure [26, 68, 71, 89]. In parallel, we additionally consider the effect of the human connectome’s geometry with a null model that approximately preserves the edge length distribution of the empirical network, in addition to its degree sequence, allowing us to strictly attribute the remaining effects to topology [18].

We initially focus on the topological shortest path (hereafter referred to as a “path”), because (a) it is a simple and fundamental method to infer communication, and (b) in many classes of networks, including brain networks, alternative communication mechanisms nevertheless take advantage of shortest paths without any knowledge of the global topology, including diffusion [44, 45, 89] and navigation [108–111, 134]. We then investigate inter-regional communication capacity, mapping it onto large-scale cognitive systems and patterns of functional specialization. Finally, we also consider the relationship between spatial proximity/geometric embedding and communication capacity, as well as alternative communication mechanisms.

## RESULTS

The results are organized as follows. First, we develop a method to standardize communication capacity between pairs of brain regions. We then relate inter-regional communication to the brain’s spatial embedding, canonical functional systems and patterns of functional specialization. All analyses were conducted in a sample of *N* = 69 healthy participants (source: Lausanne University Hospital; DOI: 10.5281/zenodo.2872624 [51]; see *Methods* for detailed procedures):

- *Structural connectivity*. Structural connectivity was reconstructed from individual participants’ diffusion spectrum imaging data using deterministic streamline tractography. A distance-dependent consensus-based thresholding procedure was then used to assemble a group-representative weighted structural connectivity matrix of streamline density [19, 88, 89].
- *Functional connectivity*. Functional connectivity was estimated from the same individuals’ restingstate functional MRI (rs-fMRI) data using pairwise Pearson correlations among regional time courses. Fisher’s r-to-z transformation was applied to individual functional connectivity matrices. A group-average functional connectivity matrix was then computed as the mean across individuals, which was back-transformed to correlation values.

The sample was randomly divided into *Discovery* (*n* = 34) and *Validation* (*n* = 35) subsets. Analyses were conducted in a high (1000 nodes) and low (219 nodes) resolution parcellation using the Cammoun atlas [24], a subdivision of the Desikan-Killiany anatomical atlas [35]. See *Sensitivity analyses* for details.

### Benchmarking dyadic communication capacity

To quantify polysynaptic communication capacity between pairs of brain regions, we first compute the topological weighted shortest path lengths on the unthresholded structural connectome (Fig. 1) [36]. Shorter weighted path length between a pair of regions indicates greater communication capacity [6, 7, 107]. We simultaneously construct a population of rewired networks that preserve the density and weighted degree sequence of the empirical network [89, 132]. We then compute the path lengths for each rewired network, indicating the communication capacity between pairs of brain regions under the null hypothesis that inter-regional relationships depend only on weighted degree and density (Fig. 1). Finally, we standardize element-wise the empirical path lengths against the population of path lengths in the rewired null networks. The resulting standardized shortest path length matrix quantifies in terms of z-scores how unexpectedly short (< 0) or unexpectedly long (> 0) communication pathways are between any given pair of brain regions.

**Figure 1.**
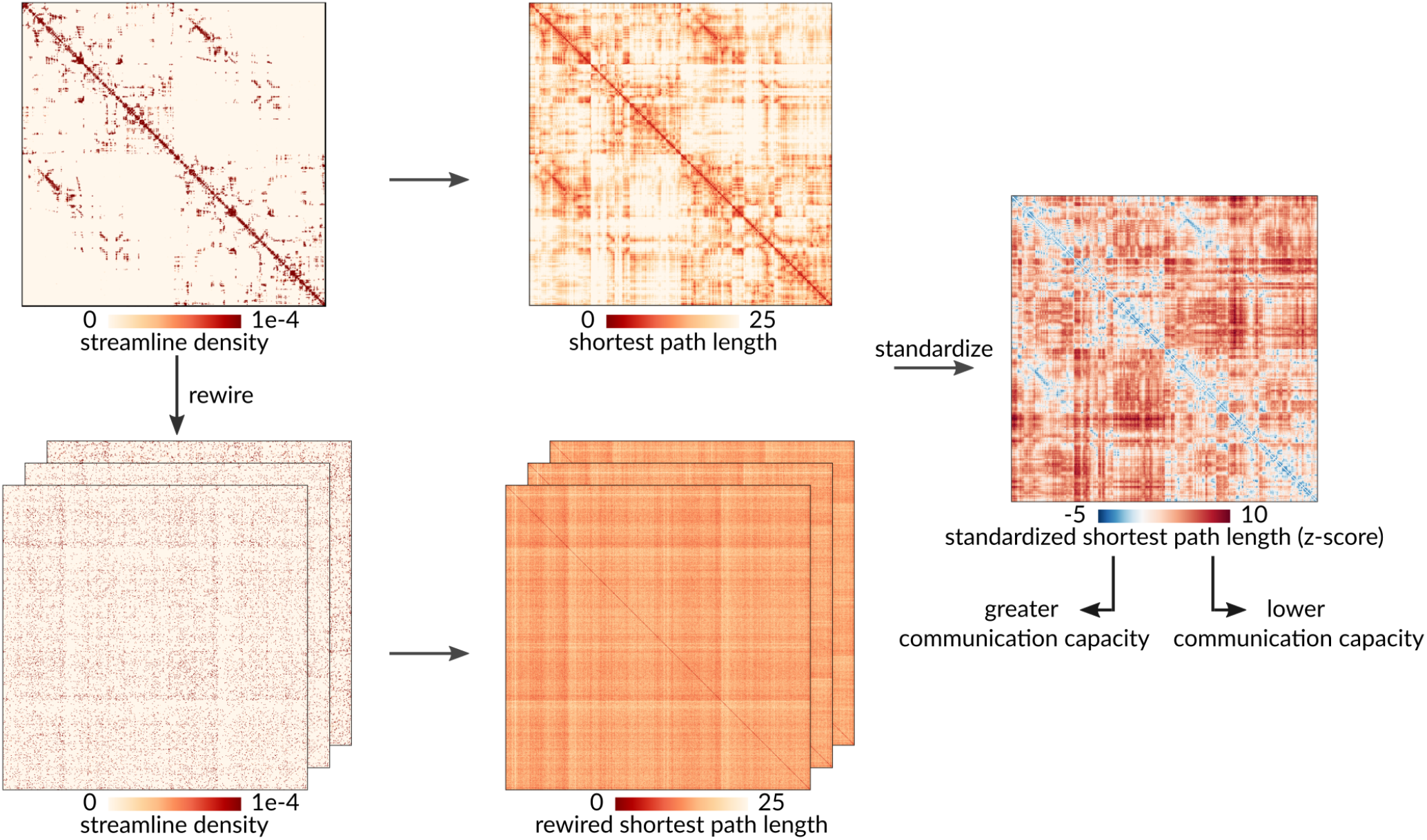
Standardization procedure. A population of null structural connectivity matrices that preserve the size, density, and weighted degree sequence of the empirical group-consensus network was generated by randomly rewiring pairs of edges. Weighted shortest path lengths were then computed between every pair of brain regions for the empirical structural brain network and each rewired null. Finally, the path lengths of the empirical network were standardized element-wise against the null population of path lengths from the rewired networks. Lower standardized shortest path length indicates greater communication capacity.

Fig. 2a shows a scatter plot between empirical (abscissa) and rewired (ordinate) path lengths, where each point represents a pair of regions. As expected, the majority of points fall below the identity line (87.15%), suggesting that most path lengths in rewired networks are shorter than in the empirical structural brain network [138]. This is in line with numerous global accounts of the shortest path length of random networks and their comparison with characteristic path lengths of empirical brain networks [2, 9, 59, 118, 138]. Interestingly, a number of points reside above identity (12.85%), suggesting that these region pairs enjoy greater-than-expected capacity for communication. Fig. 2b further demonstrates this result, showing the distribution of standardized path lengths for all pairs of regions. Negative values indicate dyads with greater-than-expected communication capacity, and positive values indicate dyads with lower-than-expected communication capacity.

**Figure 2.**
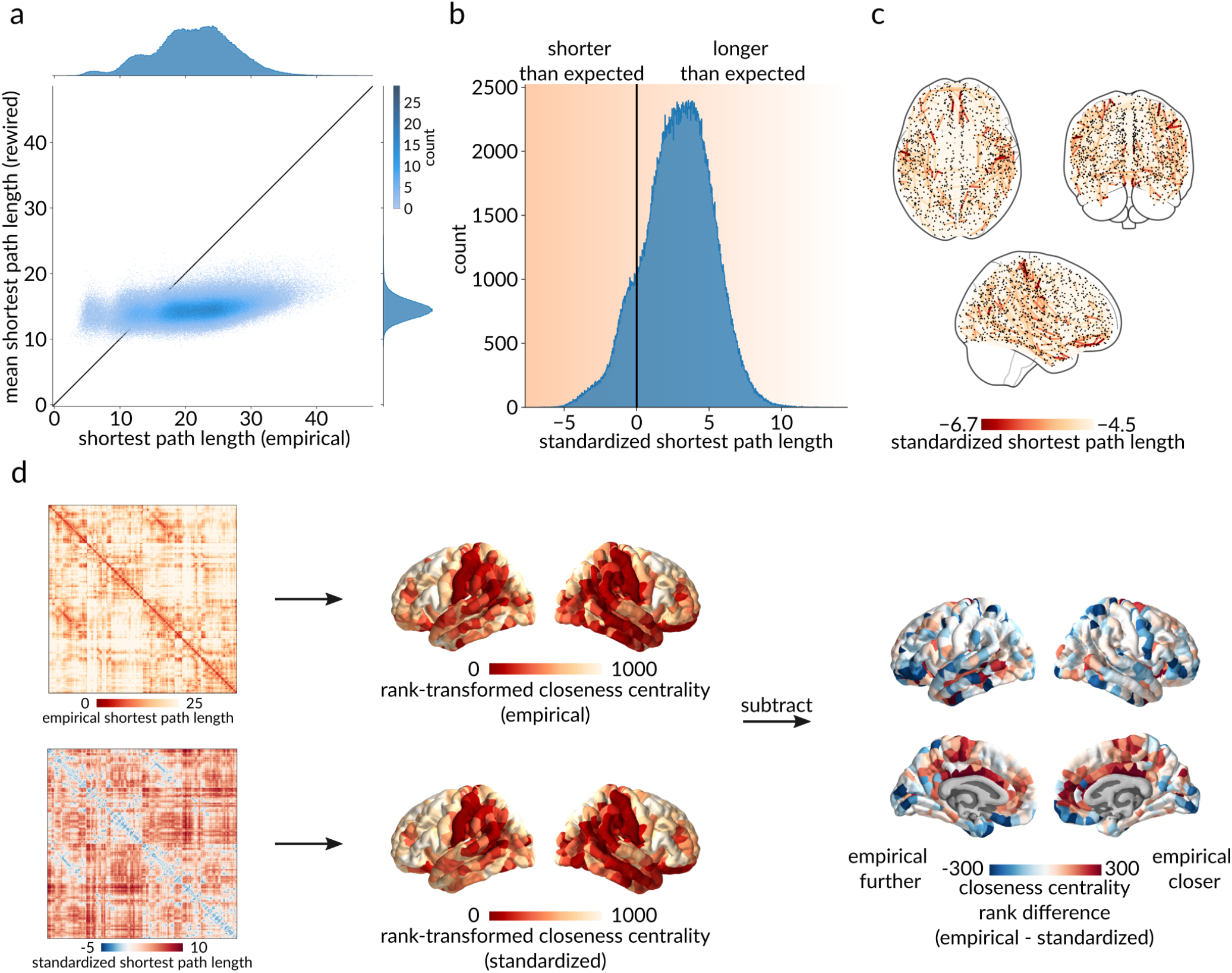
Benchmarking dyadic communication capacity. (a) Scatter plot between empirical (abscissa) and rewired (ordinate) shortest path lengths, where each point represents a pair of brain regions. Marginal distribution histograms are shown on the top and right axes. Points that appear below the identity line correspond to paths with a shorter length in the rewired networks than in the empirical network, and vice versa for points above the identity line. (b) Distribution of standardized shortest path lengths (z-scores) for all pairs of brain regions. Values less than 0 indicate greater-than-expected communication capacity, and values greater than 0 indicate lower-than-expected communication capacity. (c) Spatial distribution of the top 1% unexpectedly short path lengths (d) Using the empirical and the standardized path length matrices, closeness centrality (inverse mean path length to the rest of the network) was computed and rank-transformed for every brain region. The two resulting brain maps were then subtracted, resulting in a brain map of the region-wise differences between closeness centrality ranks in the empirical and the standardized networks. Red regions are more integrated in the empirical network, and blue regions are more integrated in the standardized network.

To get a sense of how the centrality or “hub-ness” of each brain region changes when path lengths are standardized, we compute the closeness centrality (inverse mean path length to the rest of the network) of each brain region using the empirical and the standardized path length matrices. Fig. 2d shows the difference between rank-transformed closeness computed using empirical and standardized path lengths. The figure suggests that the inferred importance of a brain region changes considerably when the procedure is applied. Namely, red regions (e.g. cingulate cortex) are more central in the empirical shortest path length network, and blue regions (e.g. orbitofrontal cortex) are more central in the standardized shortest path length network.

### Communication pathways delineate functional systems

We next consider how communication paths can be contextualized with respect to canonical features of brain networks, including spatial embedding, structurefunction coupling and macroscale intrinsic network organization. Fig. 3a shows the relationship between standardized path length and pairwise inter-regional physical distance (left) and pairwise inter-regional functional connectivity (right). There is a positive association between physical distance and standardized path length, consistent with the notion that areas that are physically further apart have lower communication capacity [105, 111, 120]. There is also a negative association between standardized path length and functional connectivity, consistent with the notion that pairs of areas that are topologically closer have more coherent time-courses [45, 60]. Collectively, these results show that standardized path length recapitulates well-known and expected relationships between the topology, geometry and functional connectivity of the brain [121].

**Figure 3.**
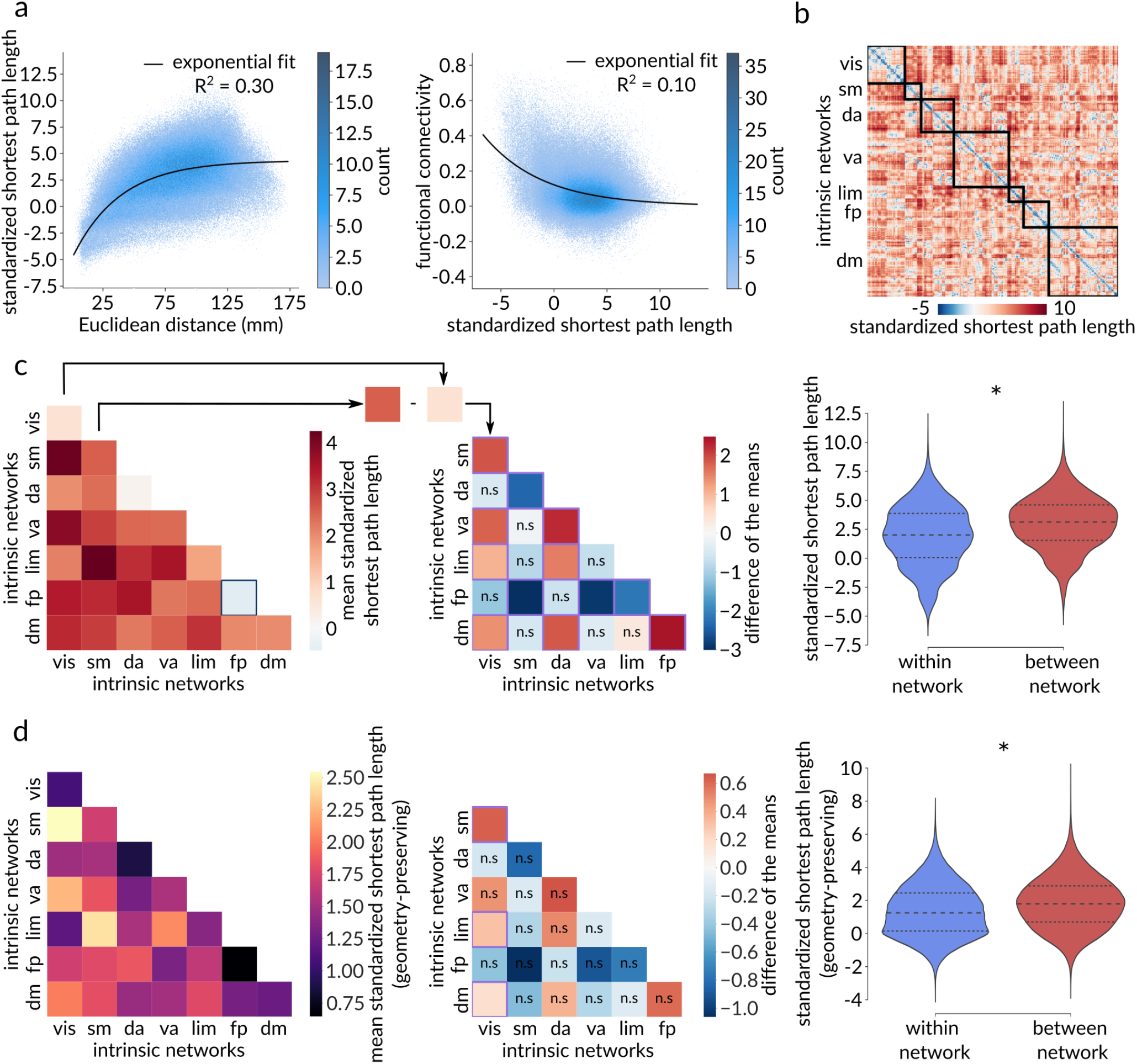
Communication pathways delineate functional systems. (a) Standardized shortest path length (***y***) between two brain regions grows as a function of the Euclidean distance (***x***) between them (left). The black line corresponds to the fitted exponential ***y*** = – **9.31*e***^-0.03*x*^ + **4.33**. Functional connectivity (***y***) between two brain regions decays as a function of the standardized shortest path length (**x**) between them (right). The black line corresponds to the fitted exponential ***y*** = **0.12e**^-0.18*x*^. (b) Standardized shortest path length matrix with brain regions ordered based on their affiliations to the Yeo intrinsic networks. (c) Left: Heatmap of the mean standardized path lengths across node pairs belonging to the same intrinsic network (diagonal) and to different intrinsic networks (off-diagonal). A blue square identifies a negative mean standardized path length, indicative of shorter-than-expected communication pathways with greater-than-expected communication capacity. Middle: Heatmap of the pairwise differences of the means among Yeo intrinsic networks, calculated as the mean value of the network on the y-axis minus the mean value of the network on the x-axis, with the mean value corresponding to the mean standardized path length across node pairs belonging to the same network (diagonal elements of the left heatmap). A purple square indicates significant difference of the means based on network label permutation using spatial autocorrelation-preserving null models (Bonferroni corrected, ***α*** = **.05**), whereas “n.s.” denotes not significant differences. The frontoparietal network displays a consistently shorter mean standardized path length (i.e., higher internal communication capacity) compared to other networks, whereas the somatomotor network exhibits a consistently greater mean standardized path length (i.e., lower internal communication capacity) in comparison to other networks. Right: The mean within-network standardized path length is significantly shorter than the mean between-network standardized path length (*p*_spin_ < .001). (d) Same as (c) but for shortest path lengths standardized using a geometry-preserving null model. Intrinsic networks: vis = visual, sm = somatomotor, da = dorsal attention, va = ventral attention, lim = limbic, fp = frontoparietal, dm = default mode.

How are communication pathways organized among the canonical macroscale intrinsic networks? Resting-state functional connectivity networks are communities of functionally related areas with coherent time-courses that are thought to be putative building blocks of higher cognition [15, 33, 102, 144], but how these networks map onto the underlying communication pathways is not completely understood [7, 121]. To address this question, we first stratify brain regions according to their membership in the intrinsic networks derived by Yeo, Krienen and colleagues [144] (Fig. 3b). Fig. 3c (left) shows the mean standardized path length within each intrinsic network (diagonal) and between all pairs of intrinsic networks (off diagonal). We generally observe shorter path lengths within networks compared to between networks; Fig. 3c (right) confirms this intuition, showing that the mean within-network path length is significantly shorter than the mean between-network path length (*p*_spin_ < .001).

Next, we quantify and compare the internal communication capacities of pairs of intrinsic networks by computing the difference between their respective within-network mean standardized path length (Fig. 3c, middle). We find that the frontoparietal network has consistently greater internal communication capacity compared to other networks, while the somatomotor network has consistently lower internal communication capacity compared to other networks. Interestingly, Fig. 3c (left) also indicates that communication pathways internal to the frontoparietal network are the only ones to exhibit a greater-than-expected communication capacity, characterized by a negative mean standardized path length.

Moreover, we reproduce this analysis using a geometry-preserving null model (Fig. 3d). Once again, we find significantly shorter path lengths within Yeo networks (*p*_spin_ < .001; Fig. 3d, right) and identify the frontoparietal network and the somatomotor network as consistently exhibiting the highest and lowest internal communication capacity, respectively (Fig. 3d, middle). However, most pairwise comparisons between intrinsic networks are now not significant as assessed by spatial autocorrelation-preserving nulls (Bonferroni corrected, ***α*** = **.05**). Furthermore, the frontoparietal network does not present a negative mean standardized path length anymore, indicating that its greater-than-expected communication capacity can be partly attributed to its spatial embedding (Fig. 3d, left).

### Communication capacity and functional specialization

Given that communication capacity is regionally heterogeneous and maps onto intrinsic networks, we ask whether regional communication capacity is related to functional specialization. Fig. 4a shows the mean standardized path length from each region to the rest of the network, with red indicating greater integration with the network and yellow indicating lower integration.

**Figure 4.**
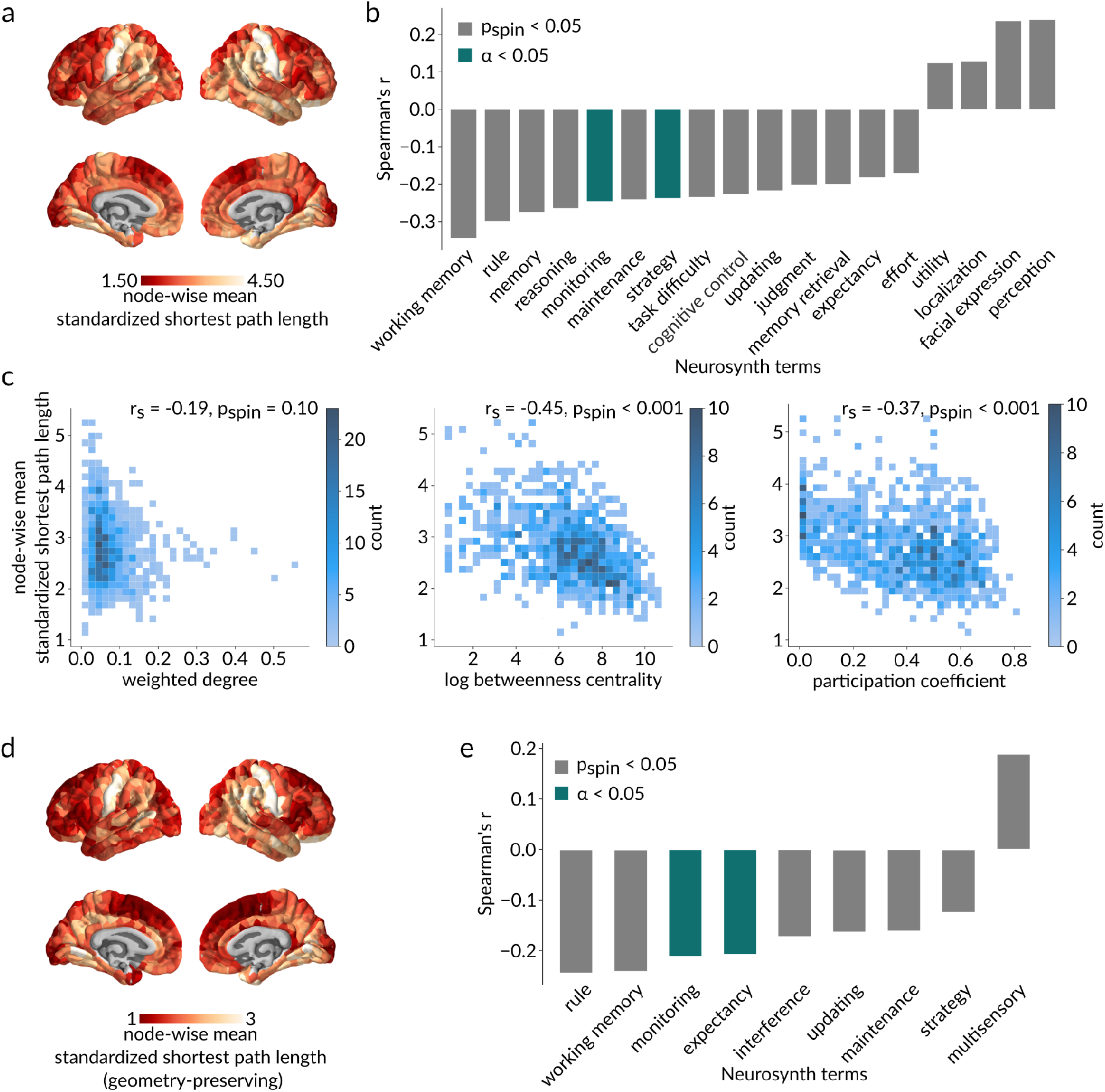
Communication capacity and functional specialization. (a) Brain map of mean standardized path length from each node to the rest of the network, with red denoting a greater integration of the node within the network and yellow denoting a lower integration. (b) Statistically significant spatial correlations based on spatial autocorrelation-preserving nulls (*p*_spin_ < .05 in grey; Bonferroni corrected, ***α*** = **.05** in green) between node-wise mean standardized path lengths and meta-analytic probabilistic functional activation maps from the Neurosynth platform, associated with 123 terms overlapping the Neurosynth database and the Cognitive atlas. An important set of anticorrelations suggests that highly integrated nodes are associated with higher-order cognitive functions. (c) Relationships between node-wise mean standardized path length and topological features of the empirical weighted structural network. As expected due to the standardization procedure, node-wise mean standardized path length is not significantly correlated with weighted degree (*r*_s_ = −.19, *p*_spin_ = .10; left), while it is significantly negatively correlated with betweenness (*r*_s_ = −.45, *p*_spin_ < .001; middle) and participation (*r*_s_ = −.37, *p*_spin_ < .001; right), suggesting that regions that are more topologically integrated also have a more diverse connection profile among functionally specialized intrinsic networks. (d-e) Same as (c) but for shortest path lengths standardized using a geometry-preserving null model.

We statistically compare this map with a library of meta-analytic task-based fMRI activation maps from the Neurosynth repository [56, 142]. Each of the Neurosynth brain maps consists of region-wise measures of the probability that a particular term is reported in a study if an activation was observed in a given region. In this analysis, we only consider the intersection of terms from the Neurosynth database and the Cognitive atlas [100], comprising a total of 123 cognitive and behavioral terms. Fig. 4b shows statistically significant spatial correlations between the node-wise standardized path length map and each of the Neurosynth term maps, as assessed using spatial autocorrelation-preserving null models [3, 82] (*p*_spin_ < .05 in grey; Bonferroni corrected, ***α*** = **.05** in green). We find anticorrelations with higher-order cognitive terms (e.g. “monitoring”, “strategy”, etc.). This suggests that areas that communicate closely with many other areas in the connectome are associated with high-order cognitive function. In other words, cognitive integration appears to be supported by network integration.

To better understand why lower standardized path length is associated with higher-order cognitive function, we compare the regional map of standardized path length with maps of weighted degree (sum of edge weights incident on a node; strength), betweenness (proportion of shortest paths that traverse a node) and participation (distribution of node links among functional network communities) (Fig. 4c). All three topological features were computed on the empirical weighted structural network. As expected due to the standardization procedure, there is no significant correlation with weighted degree (*r*_s_ = −.19, *p*_spin_ = .10). However, standardized path length is significantly negatively correlated with betweenness (*r*_s_ = −.45, *p*_spin_ < .001) and participation (*r*_s_ = −.37, *p*_spin_ < .001), suggesting that regions that are closely integrated into the connectome can better sample information from multiple specialized communities.

Furthermore, we replicate this analysis using a geometry-preserving null model (Fig. 4d-e). Once again, we find significant anticorrelations between node-wise standardized path lengths and higher-order cognitive terms (e.g., “monitoring”, “expectancy”). This suggests that the observed association between cognitive and network integration is not passively endowed by the brain’s physical embedding, but rather driven by its topological organization.

### Extending to multiple communication models

So far, we have only considered path length as a proxy for communication. However, there exist numerous other models of communication in the connectome [7]. Here we extend the framework developed in the previous sections to additional measures of communication proximity that have been proposed for brain networks, including search information ([45, 106]), path transitivity ([45]), communicability [30, 39] and mean first-passage time [44, 94]. As before, we first standardize each communication matrix using a population of rewired networks (Fig. 5a). We then extract all dyadic (*i, j*) elements from each communication matrix and assemble them into the columns of a dyads × communication models matrix.

**Figure 5.**
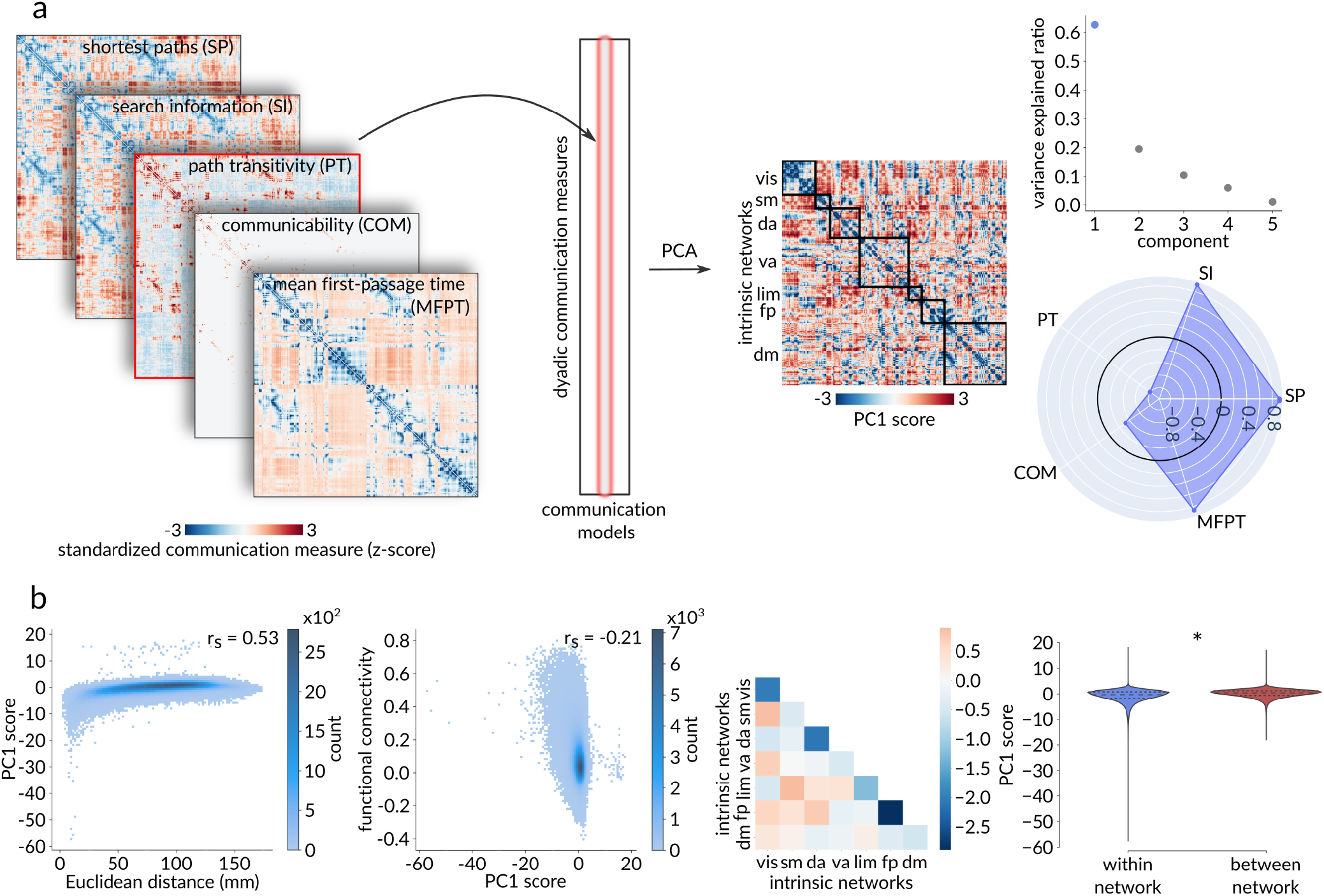
Extending to multiple communication models - dyad-level. (a) Standardized dyadic communication measures across five communication models were assembled into the columns of a dyads × communication models matrix. Principal component analysis, applied to this matrix, identified a single dominant component that accounts for 62.79% of the dyadic-level variance in communication capacity. The radar chart represents the first component’s loadings (i.e., correlations with the five communication models under consideration), with the greatest contribution to the aggregated communication measure coming from search information, path length, and mean first-passage time, with only a minor contribution from communicability. (b) The aggregate communication measure yields similar results to the standardized shortest path length. From left to right: Positive Spearman correlation between PC1 score and Euclidean distance (*r*_s_ = .53, *p* ≈ 0), negative Spearman correlation between functional connectivity and PC1 score (*r*_s_ = −.21, *p* ≈ 0), and significantly lower within-network than between-network PC1 score (*p*_spin_ < .001).

Applying principal component analysis (PCA) to this matrix identifies a single dominant component that accounts for 62.79% of the dyad-level variance in communication. The resulting PC1 scores are then used to constitute an “aggregated” communication matrix that shows the capacity for communication among all pairs of brain areas across multiple communication models. The greatest contribution to the aggregated communication measure is from search information, path length and mean first-passage time, with modest contribution from communicability. Note that, unlike the original communication measures, this aggregated communication measure is not necessarily transitive.

Overall, we observe similar results using the aggregated, multi-communication measure (Fig. 5). Namely, we find a positive relationship between standardized communication distance and physical distance (*r*_s_ = .53, *p* ≈ 0; Fig. 5b), a negative relationship between standardized communication distance and functional connectivity (*r*_s_ = −.21, *p ≈* 0; Fig. 5b), and significantly shorter communication distance within canonical intrinsic networks than between networks (*p*_spin_ < .001; Fig. 5b).

Fig. S1a shows the same procedure applied at the node level. As before, we find consistent results, showing that areas that are closer in communication distance to other areas in the connectome tend to be associated with higher-order cognitive function (Fig. S1c) and greater participation in network communities (*r*_s_ = −.40, *p*_spin_ < .001; Fig. S1d; right).

### Disentangling the contributions of topology and geometry

Physical proximity is an important predictor of connection probability and connection weight [18, 105], and therefore by extension, communication capacity between areas. We therefore seek to disentangle the contribution of geometry from the contribution of topology to the results reported so far. Initially, we utilized degree-preserving rewiring to construct populations of null networks and standardize path length [85, 89]. To additionally control for the role of geometry, we repeat the experiments using a more conservative null model that approximately preserves the edge length distribution and the edge length-to-weight mapping, in addition to degree sequence [18].

Fig. S2 shows the results when applying the geometry-preserving null model. As for the degree-preserving null model, Fig. S2a shows that the majority of path lengths are shorter in the rewired networks than in the empirical network (86.04%). However, the majority of points (region pairs) now lie closer to the identity line, suggesting that the physical embedding of the structural brain network contributes in pulling its regions apart topologically. Furthermore, controlling for geometry in the standardization procedure accentuates the integration and segregation of certain regions, providing a clearer, more homogeneous picture of changes in closeness centrality (Fig. S2b). The orbitofrontal cortex becomes more integrated and the cingulate cortex and precuneus become more segregated when strictly considering the empirical network’s topology. Interestingly, the paracentral lobule and the motor cortex now move in directions opposite to those observed when using the null model that did not preserve the connectome’s geometry. Indeed, the geometry-preserving standardization procedure moved the paracentral lobule topologically closer to the rest of the network and the motor cortex further. This suggests that the physical distance of these regions to the rest of the brain interacts with its topology to centralize the motor cortex and segregate the paracentral lobule.

As expected, controlling for edge length considerably attenuates the relationship between standardized path length and Euclidean distance (Fig. S2c, left). Importantly, most of the other results are largely preserved, including shorter standardized path lengths within compared to between networks (*p*_spin_ < .001; Fig. 3d, right) and a relationship between node-wise communication and integrative function (Fig. 4e). Collectively, this control experiment suggests that most of the results reported previously – except the relationship with Euclidean distance – are mainly driven by the topological organization of the connectome rather than spatial embedding and geometric relationships. An important exception is the higher-than-expected internal communication capacity of the frontoparietal network, which seems to have been partly driven by the brain’s geometry.

### Sensitivity analyses

We test the replicability of the findings in the *Validation* sample (Fig. S3) and seek to assess the sensitivity of the results to a variety of processing choices. We repeat all analyses using a lower parcellation resolution of 219 nodes (Fig. S4) and binary (non-weighted) structural networks (Fig. S5).

We find consistent results across all sensitivity analyses. This includes significantly shorter communication pathways between cortical regions belonging to the same intrinsic functional network than between regions belonging to different intrinsic networks, as well as significant relationships between a region’s topological integration and its association with higher order executive functions (Fig. S3, S4, S5). The frontoparietal network is consistently identified as exhibiting the highest internal communication capacity. Communication pathways internal to the frontoparietal network also have a negative mean standardized path length, indicative of a greater-than-expected communication capacity (Fig. S3, S4, S5).

Next, we seek to test the extent to which the results are influenced by the inclusion of direct monosynaptic pathways (i.e., paths between directly anatomically connected nodes; Fig. S6). We therefore repeat the analyses in strictly polysynaptic communication pathways (i.e., paths between pairs of nodes that are separated by two or more anatomical connections) (Fig. S6c, Fig. S7, Fig. S8, Fig. S9). Again, we find consistent results with the notable exception of the communication capacity of the fronto-parietal network (Fig. S7; original result in Fig. 3c). This indicates that the greater-than-expected communication capacity of this network is partly driven by monosynaptic connections.

Recently, there has been a growing interest to relate patterns of brain communication with the higher-order connectivity of brain networks ([12, 29, 52, 115]). In alignment with this new line of inquiry, and to demonstrate the flexibility of our standardization procedure, we apply it to a recently introduced higher-order multimodal communication measure: structure-function bandwidth ([95]). Based on a multilayer framework in which structural and functional connectivity are considered simultaneously, bandwidth between two brain regions in the functional connectivity layer is defined according to the minimum edge weight of a path connecting these nodes in the structural connectivity layer. The maximum bandwidth is then selected across all paths of a given length. Here, we consider triangles composed of two-hop structural paths closed by a functional edge. We weigh structure-function bandwidth by the functional edge weights to provide a generalized communication measure taking into account all functional connections enclosing triangles. This yields a structurefunction bandwidth matrix which we concurrently standardize using the same procedure previously described, with structure-function bandwidth being recomputed in a population of rewired structural networks while the functional network is left intact. We then consider how our generalized structure-function bandwidth measure is organized among intrinsic functional networks. We observe distinct patterns of intra- and inter-network structure-function bandwidth when comparing the empirical and the standardized measures (Fig. S10). This analysis demonstrates how the standardization procedure can be readily extended to accommodate questions about higher-order brain network architecture.

## DISCUSSION

In the present report, we introduce a simple method to standardize communication path lengths in brain networks. These results showcase how dyadic relationships can be resolved and studied while accounting for more basic topological and geometric features of the network. Building on hierarchically constrained null models, this rigorous standardization procedure enables specific quantification and localization of the degree of unexpectedness of communication measures. In contrast to classical approaches, considering communication capacity at a finer granularity allows us to revisit previous investigations of brain hubs, recapitulate well-known geometric and functional attributes of inter-regional communication, and uncover new relationships between the human connectome’s topological and functional architectures across multiple topological levels of analysis in a nuanced and principled way.

Classical studies focused on global path length or efficiency of brain networks. These reports found evidence of near-minimal path length characteristic of small-world architecture across multiple species and reconstruction methods [11, 59, 67, 118, 138]. In addition, empirical studies found that low characteristic path length or high global efficiency is associated with greater cognitive performance [78, 130] and is concomitant with healthy neurodevelopment [13, 40, 55, 69, 70]. Altogether, these findings speak to the behavioral and biological relevance of global accounts of communication capacity.

However, global communication measures such as characteristic path length or global efficiency are effectively summary statistics of a myriad of complex interregional communication relationships. In the present study, we focus specifically on dyadic communication while statistically controlling for fundamental topological and geometric features (i.e. degree and spatial position). We find that most communication pathways between brain regions are longer than expected on the basis of their degree and/or spatial position. Despite the standardization procedure, we still recapitulate fundamental features of inter-regional communication, such as positive relationships with spatial proximity [105, 111, 120] and negative relationships with restingstate functional connectivity [45, 60, 110, 133, 145].

We note that the procedure used here is similar to what is typically done when considering global communication statistics, such as characteristic path length. Namely, characteristic path length is often normalized by the mean characteristic path length across a population of null networks, such as when estimating the small-world coefficient [61, 138]. We build on this general approach by finely resolving dyadic communication relationships. In addition, by standardizing each dyadic path length as opposed to normalizing by the mean, we implicitly take into account variance across the null population [10].

The standardization procedure alters the centrality rankings of brain regions, suggesting that taking the constraints of degree into account can lead to different inferences about the functional importance of brain regions. For instance, we find that the orbitofrontal cortex is more integrated in the standardized network than the empirical network, and that parts of the medial prefrontal cortex, cingulate cortex, and precuneus are less integrated in the standardized network. These results run counter to numerous classical investigations of brain hubs that did not explicitly control for degree when estimating different centrality measures based on path length, such as betweenness and closeness [54, 127, 129].

A large body of neuroimaging studies has highlighted the existence of a number of macro-scale communities of functionally related brain regions with correlated resting-state fMRI signals [33, 102, 123, 144]. Previous efforts were made to relate these patterns of synchronized spontaneous activity to the underlying anatomical scaffolding of the brain [126]. Specifically, some studies have investigated the structural underpinnings of the functional links within and between resting-state networks by identifying specific white matter tracts that could mediate these relationships [50, 125]. More recently, a growing interest in the functional predictive utility of communication measures based on structural connectivity has led to investigations into the relationships between functional brain networks and underlying patterns of polysynaptic communication. It has been shown that the intrinsic functional hierarchy of the brain guides communication trajectories and allows for signal diversification [134]. Furthermore, it was found that the modular boundaries of resting-state functional connectivity were approximated by modules of polysynaptic communication distance [109]. In line with these studies, we map communication pathways among canonical intrinsic functional networks [144]. We find that standardized path lengths are significantly shorter within than between intrinsic networks, suggesting that the topological organization of the human connectome contributes in giving rise to macro-scale intrinsic patterns of functional interactions.

Moreover, by organizing communication pathways within individual intrinsic networks and between pairs of networks, we identify the frontoparietal network as exhibiting the highest internal communication capacity. Previous findings had associated a greater structural global efficiency of a frontoparietal network (mean closeness centrality across nodes of the network to all brain regions) with a higher working memory capacity [99]. The present result suggests that, in addition to global integration of frontoparietal nodes, the higher-order executive control functions that have been widely attributed to frontoparietal networks [37, 41, 74, 93, 103, 135, 143], might also benefit from an unexpectedly high level of internal communication capacity.

What are the functional consequences of lower or greater communication capacity? Comparison with meta-analytic maps of functional specialization suggests that regions that are topologically closer to others tend to be associated with higher-order cognitive functions such as monitoring and strategy. In other words, we find that greater network integration is associated with cognitive integration [28, 89, 117, 141]. This is consistent with numerous theories which posit that patterns of regional specialization arise from connectivity profiles [22, 83, 84, 86, 96] and topological embedding of brain regions [98, 116, 117, 140]. Moreover, we find that nodal topological integration is positively associated with the number of shortest paths traversing it and the diversity of a region’s connections among intrinsic functional communities. This is in line with previous accounts of hub characteristics [54, 124, 127–129]. Here, we further show these features to be related even when controlling for the effects of degree and strength. Previous reports have also shown positive associations between a node’s involvement in complex tasks and the diversity and flexibility of its functional links to the rest of the brain, especially in frontoparietal regions [16, 27, 102]. Altogether, the present results complement these findings, suggesting that brain regions that subserve higher-order cognition also benefit from a structural substrate for the diversification and integration of information.

This work is part of a wider trend in the field to infer and quantify the potential for communication among brain regions based on their wiring patterns [7, 47, 48, 119, 147]. Although we mainly focus on shortest paths, multiple alternative communication protocols have been proposed [6, 17, 30, 45, 89, 91, 111, 147]. The present standardization procedure can be readily applied to any dyadic communication measure. Combining additional measures of decentralized communication such as search information and path transitivity, we find results consistent with those derived using path length. As the field moves towards more biologically realistic and validated communication protocols, future studies could adapt this standardization procedure to accommodate emerging measures of communication capacity. As an example, we show that it can be readily adapted to a higher-order multimodal communication measure defined in a multiplex brain network combining structural and functional connectivity.

In addition, the present procedure standardizes communication measures using two common types of rewiring null models. Here we focus on disentangling the contribution of the structural brain network’s topology from the background effect of spatial embedding. We therefore apply one null model that preserves the (weighted) degree sequence and another that additionally preserves wiring length [18]. Interestingly, most effects are preserved when applying the geometric null model, suggesting that they are driven by topology rather than spatial embedding. More generally, this highlights the fact that any null model – embodying a specific null hypothesis – could be used for the standardization procedure to selectively tease apart features that shape communication in brain networks [132, 146].

The present findings should be viewed in light of multiple methodological limitations. First, all networks were reconstructed using diffusion-weighted MRI, a method known to be subject to false positives and false negatives [34, 80, 122]. Although we attempt to mitigate this limitation by splitting the sample and repeating all analyses, future work is necessary to improve the quality of connectome reconstruction. Second, networks reconstructed from diffusion-weighted MRI are undirected, limiting the biological plausibility of these networks and our capacity to fit communication models to them. Third, we focus on several well-studied and mathematically fundamental communication protocols, including centralized (e.g., shortest paths) and decentralized (e.g., search information) measures, but this selection is non-exhaustive and alternative communication measures could be considered in future work.

In summary, we present a simple method to resolve communication capacity in brain networks. The method is based on conventional procedures already commonplace in connectomics, but allows researchers to focus on dyadic communication. This procedure is inherently flexible, being able to accommodate emerging communication measures and null models. As the field develops increasingly sophisticated and biologically realistic generative models of inter-regional communication, this procedure will allow greater insight into features that shape signaling patterns in brain networks.

## METHODS

Code and data used to perform the analyses can be found at https://github.com/fmilisav/milisav_dyadic_communication.

### Data Acquisition

A total of n = 69 healthy participants (25 females, age 28.8 ± 8.9 years old) were scanned at the Lausanne University Hospital in a 3-Tesla MRI Scanner (Trio, Siemens Medical, Germany) using a 32-channel headcoil [51]. The protocol included (1) a magnetization-prepared rapid acquisition gradient echo (MPRAGE) sequence sensitive to white/gray matter contrast (1 mm in-plane resolution, 1.2 mm slice thickness), (2) a diffusion spectrum imaging (DSI) sequence (128 diffusion-weighted volumes and a single b0 volume, maximum b-value 8000 s/mm^2^, 2.2 × 2.2 × 3.0 mm voxel size), and (3) a gradient echo-planar imaging (EPI) sequence sensitive to blood-oxygen-level-dependent (BOLD) contrast (3.3 mm in-plane resolution and slice thickness with a 0.3 mm gap, TR 1920 ms, resulting in 280 images per participant). The last sequence was used as part of an eyes-open resting-state fMRI (rs-fMRI) scan in which the participants were not overtly engaged in a task. Informed written consent was obtained for all participants in accordance with institutional guidelines and the protocol was approved by the Ethics Committee of Clinical Research of the Faculty of Biology and Medicine, University of Lausanne, Switzerland.

### Network reconstruction

Structural connectomes were reconstructed for individual participants using deterministic streamline tractography and divided according to a grey matter parcellation of 1000 cortical nodes [24]. The analyses were also repeated at a coarser 219 cortical regions resolution. White matter and grey matter were segmented from the MPRAGE volumes using the FreeSurfer version 5.0.0 open-source package, whereas DSI data preprocessing was implemented with tools from the Connectome Mapper open-source software [32], initiating 32 streamline propagations per diffusion direction for each white matter voxel [139]. Structural connectivity was defined as streamline density between node pairs, i.e., the number of streamlines between two regions normalized by the mean length of the streamlines and the mean surface area of the regions [54]. fMRI data underwent regression of physiological variables, including white matter, cerebrospinal fluid, and motion (estimated via rigid body co-registration). BOLD time-series were subsequently subjected to a lowpass temporal Gaussian filter with 1.92 s full width half maximum and motion “scrubbing” [101] was performed after excluding the first four time points for the time series to stabilize. Functional connectivity was then computed as the Pearson correlation coefficient between the fMRI BOLD time series of each node pair.

The data were randomly split into *Discovery* (n = 34) and *Validation* (n = 35) subsets. We then generated group-representative brain networks for each subset to amplify signal-to-noise ratio using functions from the netneurotools open-source package (https://netneurotools.readthedocs.io/en/latest/index.html). A consensus approach that preserves (a) the mean density across participants and (b) the participant-level edge length distribution was adopted for the structural connectomes [19]. First, the cumulative edge length distribution across individual participants’ structural connectivity matrices is divided into M bins, M corresponding to the average number of edges across participants. The edge occurring most frequently across participants is then selected within each bin, breaking ties by selecting the higher weighted edge on average. This procedure was applied separately for intra and inter-hemispheric edges to ensure that the latter are not under-represented. The selected edges constitute the distance-dependent group-consensus structural brain network. Finally, the weight of each edge is computed as the mean across participants. The group-representative functional connectivity matrix was defined as the group average following Fisher’s r-to-z transformation. The final group consensus matrix was back-transformed to correlation values.

### Communication models

In this section, we define the analytic communication measures associated with the network communication models considered in the present study and provide their implementation details. All the communication measures, with the exception of path transitivity, were computed using functions from the open-access Python version of the Brain Connectivity Toolbox (https://github.com/aestrivex/bctpy) [107]. Path transitivity was implemented in Python based on a MATLAB script openly available in the Brain Connectivity Toolbox.

#### Shortest paths

Let *A* denote a weighted adjacency matrix. To identify the sequence of unique edges *π_u→v_* = {*A_ui_,…, A_jv_*} spanning the minimum length between nodes *u* and *v* (i.e., shortest path), we first defined a monotonic transformation from edge weights, namely streamline density in the present case, to edge lengths *L*, which can be more intuitively considered as the cost of traversing this edge. We used the negative natural logarithm: *L_ui_* = – log(*A_ui_*), mapping greater streamline density to lower cost of signal propagation. Dijkstra’s algorithm [36] was used to identify shortest paths and their length was computed as the sum *L_uv_* = *L_ui_* + … + *L_jv_* of traversed edges’ lengths.

#### Search information

Search information quantifies the amount of information required by a naïve random walker to travel along a specific path in a network ([45, 106]). More specifically, here we consider shortest paths *π_u→v_*, capturing the accessibility of these optimal routes in network topology. This analytic measure is derived from the probability that a random walker starting at *u* follows Ω*_u→v_* = {*u*,…, *v*}, the sequence of nodes visited along *π_u→v_* to reach *v*. This probability depends on the strength of the nodes comprised in Ω*_u→v_* and can be expressed as follows:

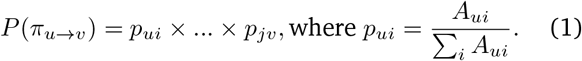

Search information can then be defined as:

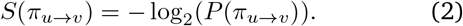

Note that this definition assumes that the random walker has no memory of its previous step, which would reduce the information needed to determine the next. Importantly, this measure is not symmetric. That is, *S*(*π_u→v_*) ≠ *S*(*π_v→u_*). Search information is contingent on the assignment of the path’s source and target nodes.

#### Path transitivity

Path transitivity is a measure of transitivity based on the shortest path between a pair of nodes. It quantifies the density of one-step detours (triangles) available along the path [45]. Intuitively, it can be considered as the accessibility of the shortest path or its robustness to a deviation of the signal traversing it. Path transitivity can be computed as the average pairwise matching index of the nodes comprising the path.

The matching index between two nodes is a measure of the similarity of their connectivity profiles, excluding edges incident on both nodes [58]. The matching index *M* between nodes *s* and *t* is defined as:

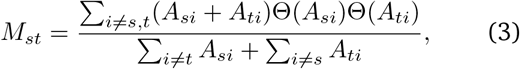

where Θ is the Heaviside step function.

Building on this definition, the path transitivity *P* of the shortest path *π_u→v_* can be defined as:

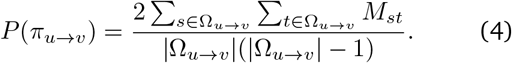

#### Communicability

Communicability between two nodes is defined as the length-weighted sum of all walks between them, with longer walks being more penalized [39]. The communicability matrix *C* of pairwise communicability estimates between all nodes in the network is calculated as the matrix exponential of the adjacency matrix: *C* = *e^A^*. Following [30], in the case of a weighted structural connectivity matrix, we first normalize the adjacency matrix as *D*^1/2^*AD*^1/2^, where *D* is the diagonal weighted degree matrix. The objective of this procedure is to reduce the undue influence of nodes with large strength.

#### Mean first-passage time

The mean first-passage time from node *u* to node *v* is the expected number of hops in a random walk evolved by a random walker starting at node *u* before arriving for the first time at node *v* [94]. Considering nodes of an undirected connected network as the states of an ergodic Markov chain allowed [44] to derive the mean first-passage time *T* between nodes *u* and *v* as:

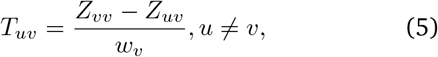

where *w* is the steady state vector of the underlying Markov process and *Z* is the fundamental matrix calculated as (*I* – *P* + *W*)^-1^. *P* is the transition probability matrix computed as D^-1^*A*, where *D* is the diagonal weighted degree matrix. *W* is a square matrix whose columns correspond to w. In the present study, following [145], we standardize as z-scores the columns of the matrix of pairwise mean first-passage time among all nodes in the network to remove nodal bias.

#### Structure-function bandwidth

Structure-function bandwidth quantifies how well structural connectivity throughput mediates functional connectivity in a multiplex network composed of a structural connectivity layer and a functional connectivity layer. Bandwidth between two nodes in the functional connectivity layer is defined according to the minimum edge weight of a path connecting these nodes in the structural connectivity layer. The maximum bandwidth is then selected across all paths of a given length. Here, we consider triangles composed of two-hop structural paths closed by a functional edge. We weigh structure-function bandwidths by the functional edge weights to provide a generalized communication measure taking into account all functional connections enclosing triangles. The functional connectivity-weighted bandwidth *B* between nodes *u* and *v* is computed as follows:

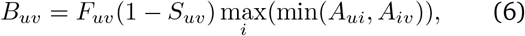

where *F* is the weighted functional connectivity matrix, *S* is the binary structural connectivity matrix. The (1 – *S_uv_*) term excludes all cases where a direct structural edge also connects nodes *u* and *v*. Every possible intermediary node *i* in the structural connectivity layer is considered to complete the triangle.

### Null models

#### Standardization

Network features are interrelated, with many complex network properties depending on basic features [132]. To mitigate the effect of differences in simple features on the topological relationships under study, we provide a frame of reference to communication measures. First, we built a population of 100 null structural connectivity matrices by randomly rewiring pairs of edges of the empirical group-consensus networks, systematically disrupting their topology while maintaining basic network features. Next, communication measures were computed for every node pair of the empirical structural brain networks and all 100 surrogate graphs. Finally, the empirical communication matrices were demeaned and standardized to unit variance elementwise using the null population. This procedure yielded, in units of standard deviation, an approximate communication measure in relation to what would be expected by chance in a similar network.

Two null models were considered as part of this procedure, resulting in a hierarchy of preserved network features. Both surrogate models guarantee the connectedness of the produced rewired network, that is, no node is disconnected from the rest of the network.

The first method randomly swaps pairs of edges (approximately 10 swaps per edge) while maintaining network size (i.e., number of nodes), density (i.e., proportion of possible edges expressed), and degree sequence (i.e., number of edges incident to each node) [85]. We applied an implementation of this technique openly available in the Python version of the Brain Connectivity Toolbox (https://github.com/aestrivex/bctpy) [107]. To extend this procedure to weighted structural connectivity matrices, we then used simulated annealing to further preserve the empirical network’s strength sequence (sum of edge weights incident to each node) [89]. Simulated annealing is a stochastic search method used to approximate the global optimum of a given function [71]. This is achieved through the Metropolis procedure [87], controlled by the temperature parameter *T*. The simulated annealing process is initiated at a high temperature which allows the exploration of costly system configurations, preventing the algorithm from getting stuck in local minima. Throughout the process, the system is slowly cooled down while descending along the optimization landscape, with increasingly limited uphill rearrangements and smaller, fine-tuned changes in the system cost. Here, we minimize the cost function *E* defined as the sum of squares between the strength sequence vectors of the empirical and the randomized networks: *E* = ∑*_i_* (*s_i_′* – *s_i_*)^2^, where *s_i_* and *s_i′_* are the strengths of node *i* in the empirical and the null networks, respectively. To optimize this function, edge weight pairs were randomly swapped. Rearrangements were only accepted if they lowered the cost of the system or met the probabilistic Metropolis acceptance criterion: *r* < exp(-(*E′* – *E*)/*T*), where *r* ~ *U*(0, 1). We used an annealing schedule composed of 100 stages of 10000 permutations with an initial temperature of 1000, halved after each stage.

A limitation of this rewiring procedure is that, on average, randomly flipping pairs of edges will result in unrealistically large distances between nodes [132]. To address this issue, we use a Python implementation of an approach proposed in [18], openly available as a function in the netneuro-tools package (https://netneurotools.readthedocs.io/en/latest/index.html). This method applies the same iterative rewiring procedure as the first, but with additional constraints. Edges are binned according to the Euclidean distance between the centroids of their associated parcel pair. The number of bins was determined heuristically as the square root of the halved number of edges in the empirical group-consensus network. Swapping is then performed within each bin to approximately preserve the edge length distribution, in addition to exactly reproducing the network’s size, density, and degree sequence like the first method. The maximum total number of edge swaps to perform was set to the network size times 20. In maintaining the geometry of the empirical network, this rewiring model provides a more representative surrogate, resulting in a more conservative null model.

Collectively, the two null models used for the standardization of the structural connectivity matrices express a hierarchy of constraints. When considered in parallel, they allow us to distinguish the contribution of the structural brain network’s topology from effects passively endowed by its spatial embedding when studying the connectome’s architecture [105].

#### Significance testing

To assess the significance of statistics based on node-level communication measures, we relied on spatial permutation null models [82]. We generated null distributions of statistical estimates derived from permuted brain maps, while preserving the spatial autocorrelation of the original data to respect the assumption of exchangeability.

First, we parcellated the FreeSurfer *fsaverage* surface according to the Cammoun atlas [24] using tools from the Connectome Mapper [32]. A spherical projection of the *fsaverage* surface was then used to assign spherical coordinates to each parcel; centroids were defined as the vertices closest (in terms of Euclidean distance) to the center-of-mass (i.e., arithmetic mean across all the vertices’ coordinates) of each parcel. Random rotations were then applied to one hemisphere of this spherical representation of the atlas to disrupt its topography. Rotations were then mirrored to the other hemisphere. Finally, each parcel in the original brain map was reassigned a rotated parcel using the Hungarian algorithm [73]. In contrast to other reassignment heuristics, this method attempts to reassign each rotated parcel to a unique original parcel, i.e., to retain the exact original data distribution. This is particularly important when testing network-based statistics as in the present study [82]. Overall, this procedure was repeated 1000 times using an openly accessible function from the netneurotools package (https://netneurotools.readthedocs.io/en/latest/index.html) to generate null statistical distributions.

For the correlational analyses, brain maps of communication measures were subjected to this spatial autocorrelation-preserving permutation procedure. For each 1000 rotated nulls, a correlation coefficient was computed between the surrogate brain map and the statistical map under study (i.e., maps of topological features or Neurosynth functional activation maps), yielding a null distribution of 1000 coefficients. The original rho was then compared against this null distribution to assess its significance by computing a two-sided p-value (*p*_spin_) as the proportion of more extreme null coefficients.

For the partition of the communication measures within and between Yeo intrinsic networks [144], network labels were first permuted according to the spatially-constrained Hungarian method [82]. Dyadic communication measures associated with a node pair belonging to the same intrinsic network, as identified by the permuted labels, were considered within-network, whereas measures associated with a node pair belonging to different intrinsic networks were considered between-networks. The difference between the mean values of the distributions associated with these two categories were then computed for each 1000 permutations, once again generating a null distribution of this statistical estimate against which the empirical difference was compared to produce a two-sided p-value.

A similar method was employed to assess the significance of the differences between mean internal standardized path lengths for pairs of Yeo intrinsic networks.

### Yeo intrinsic networks

When stratifying brain regions according to their membership in canonical macroscale functional systems, we used the seven intrinsic networks derived by Yeo, Krienen and colleagues [144] via clustering of resting-state fMRI data from 1000 subjects. A parcellation of the seven resting-state networks in the FreeSurfer *fsaverage5* surface space was downloaded from https://github.com/ThomasYeoLab/CBIG/. Nodes of the Cammoun parcellations were then labeled using a winner-take-all approach in which each parcel was attributed the most common intrinsic network assignment across its vertices.

### Neurosynth

Functional activation maps synthesizing more than 15000 published fMRI studies into probabilistic measures of the association between individual voxels and cognitive terms of interest were obtained from the Neurosynth platform (https://github.com/neurosynth/neurosynth [142]). This association measure quantifies the probability that a particular term is reported in a study if an activation was observed in a given region. The probabilistic maps were extracted from Neurosynth using 123 cognitive terms overlapping the set of keywords from the Neurosynth database and the Cognitive atlas [100], a public knowledge base of cognitive science. The list of cognitive and behavioral terms ranges from generic concepts (e.g., “attention”, “emotion”) to specific cognitive processes (“visual attention”, “episodic memory”), behaviours (“eating”, “sleep”), and emotions (“fear”, “anxiety”).

For each of the 123 terms, volumetric reverse inference maps were generated from Neurosynth and projected to the FreeSurfer *fsaverage5* mid-grey surface with nearest neighbor interpolation using Freesurfer’s *mri_vol2surf* function (v6.0.0; http://surfer.nmr.mgh.harvard.edu/). The resulting surface maps were then parcellated according the both the 219 and 1000 cortical nodes resolutions of the Cammoun atlas [24] to obtain node-wise mean probabilistic measures.

### Topological features

In this section, we define the topological features examined in this study and provide their implementation details. All the graph measures, with the exception of closeness centrality, were computed using functions from the open-access Python version of the Brain Connectivity Toolbox (https://github.com/aestrivex/bctpy) [107].

- *Degree*. Binary degree corresponds to the number of edges incident on a node, whereas weighted degree (strength) corresponds to the sum of edge weights incident on a node.
- *Closeness centrality*. The closeness centrality of a node is the inverse of the mean shortest path length between this node and all the other nodes in the network [107]. The closeness centrality *C* of a node *u* is defined as:

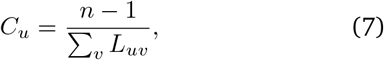

where *n* is the number of nodes in the network and *L_uv_* corresponds to the shortest path length between nodes *u* and *v*. Note that this measure can be problematic when standardizing shortest path lengths, because of negative contributions to the mean. Nevertheless, we maintain this conventional definition, because all node-wise mean standardized shortest path lengths obtained were positive.
- *Betweenness centrality*. The betweenness centrality of a node is the fraction of shortest paths between all pairs of nodes in the network that contain this node [42]. The betweenness centrality *B* of a node *i* is defined as:

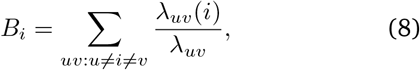

where *λ_uv_* is the total number of shortest paths from node *u* to node *v* and *λ_uv_*(*i*) is the number of these paths that include node i. Brandes’ algorithm [21] was used to compute the betweenness centrality of each individual node of the structural connectivity matrices under study.
- *Participation coefficient*. The participation coefficient is a measure of the diversity of a node’s connection profile among the communities of the network [53]. The participation coefficient *P* of a node *i* is defined as:

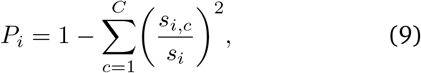

where *s_i,c_* corresponds to the sum of the weights of all edges incident on node *i* and nodes in community *c, s_i_* is the strength of node i, and *C* represents the number of communities. A participation coefficient of 0 indicates that the totality of a node’s edges are connected to other nodes within its community. The closer the participation coefficient is to 1, the more evenly distributed are its edges among the communities. Note that this measure presumes an established community structure. Here, we used the functional networks derived by Yeo, Krienen and colleagues [144] as communities to compute participation coefficients for each individual node of the structural connectomes under study.

### Principal component analysis

To take into account a range of models of communication in the brain in addition to shortest path routing, we generated an aggregate measure of the capacity for communication using principal component analysis. First, the standardized communication matrices derived following the procedures detailed above were z-scored across all elements. For each communication matrix, we then extracted all pairwise measures of communication distance, with the exception of undefined diagonal entries, and assembled them into the columns of a matrix *X* of size *d × c*, where d is the number of dyads and *c* is the number of communication models. PCA was then applied to this matrix using the *PCA* function from the scikit-learn package for machine learning in Python [97]. First, *X* was mean centered, i.e., demeaned column-wise to obtain *X_c_*. Then, a full singular value decomposition (SVD) was applied to *X_c_* such that:

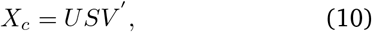

where *U* and *V* are orthonormal matrices of sizes *d × c* and *c × c*, consisting of the left and right singular vectors, respectively, and *S* is a diagonal matrix of singular values of size *c × c* [38].

In this analysis, we only kept the first component, corresponding to U_1_, the first column of *U*, which accounted for 62.79% of the total variance in dyadic communication capacity across models. The matrix multiplication *U*_1_*S* yielded the principal component scores used in this study.

The same analysis was then reproduced at the node level with vectors of standardized node-wise mean communication distance directly constituting the columns of *X*. Note that before computing the row averages of the communication matrices, we enforced symmetry of the search information and mean first-passage time matrices by replacing them with (*C* + *C^T^*)/2, where *C* is the communication matrix and *C^T^* is its transpose. In doing so, we consider both the incoming and outgoing nodal communication capacity. The same procedure was also applied to the matrix of principle component scores prior to the partition of its elements within and between Yeo intrinsic networks.

## ACKNOWLEDGMENTS

We thank Mark Nelson, Justine Hansen, Zhen-Qi Liu, Andrea Luppi, Golia Shafiei, and Estefany Suarez for helpful comments. FM acknowledges support from the Fonds de Recherche du Québec - Nature et Technologies (FRQNT). BM acknowledges support from the Natural Sciences and Engineering Research Council of Canada (NSERC), Canadian Institutes of Health Research (CIHR), Brain Canada Foundation Future Leaders Fund, the Canada Research Chairs Program, the Michael J. Fox Foundation, and the Healthy Brains for Healthy Lives initiative.

---

**Figure S1.**
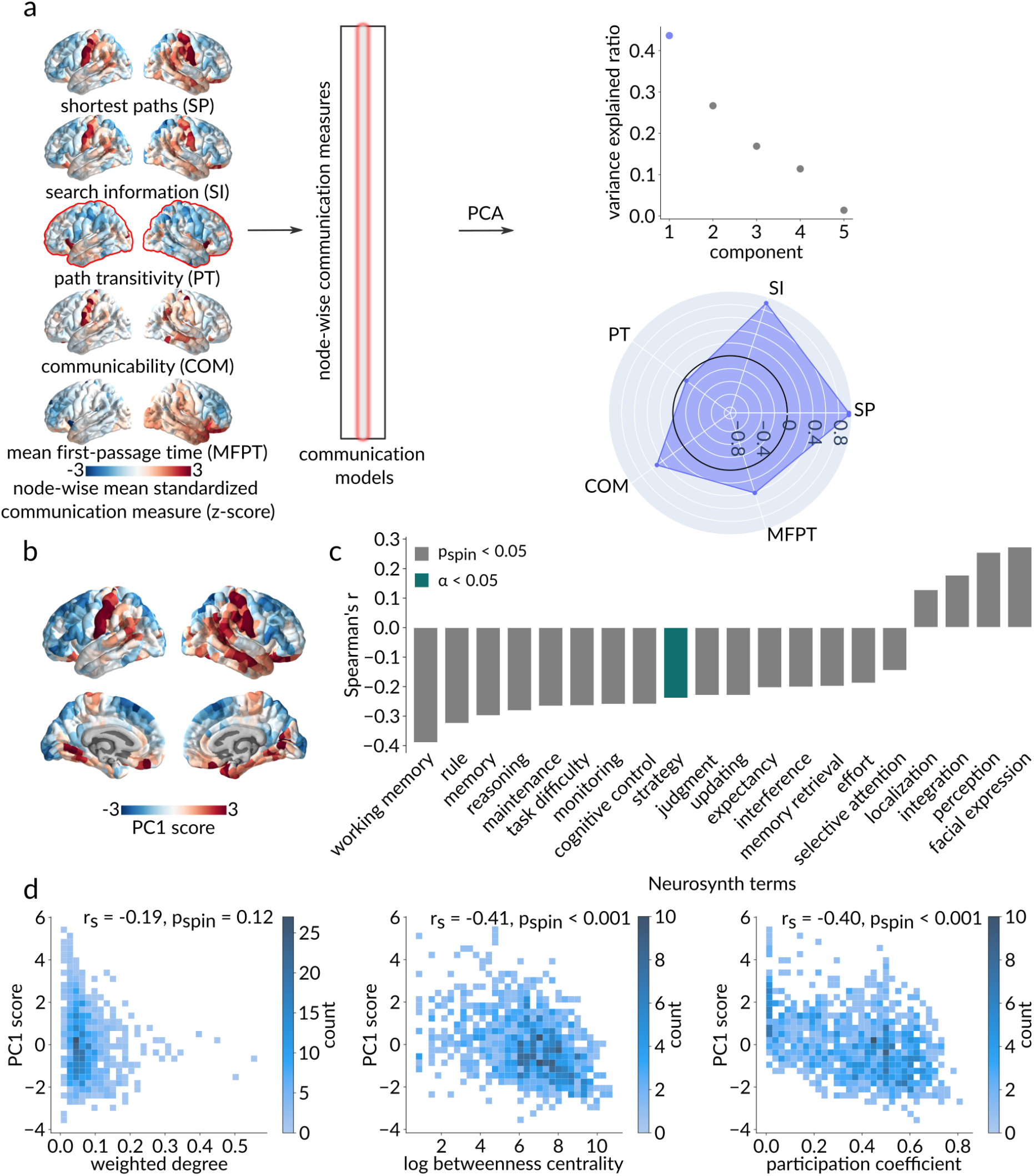
Extending to multiple communication models - node-level. (a) Standardized node-wise communication measures across five communication models were assembled into the columns of a nodes × communication models matrix. Principal component analysis, applied to this matrix, identified a single dominant component that accounts for 43.65% of the node-level variance in communication capacity. The radar chart represents the first component’s loadings (i.e., correlations with the five communication models under consideration), with the greatest contribution to the aggregated communication measure coming from search information and path length, with only a minor contribution from path transitivity. (b) Brain map of PC1 scores (c) Significant anticorrelations (*p*_spin_ < .05 in grey; Bonferroni corrected, ***α*** = **.05** in green) between node-wise mean standardized path lengths and Neurosynth functional activation maps associated to higher-order cognitive functions were replicated using PC1 scores. (d) Relationships between PC1 scores and topological features of the empirical weighted structural network recapitulate results obtained using node-wise standardized path lengths. No significant relationship was found with weighted degree (*r*_s_ = −.19, *p*_spin_ = .12; left), while a significant negative Spearman correlation was found between PC1 score and betweenness (*r*_s_ = −.41, *p*_spin_ < .001; middle) and participation (*r*_s_ = −.40, *p*_spin_ < .001; right).

**Figure S2.**
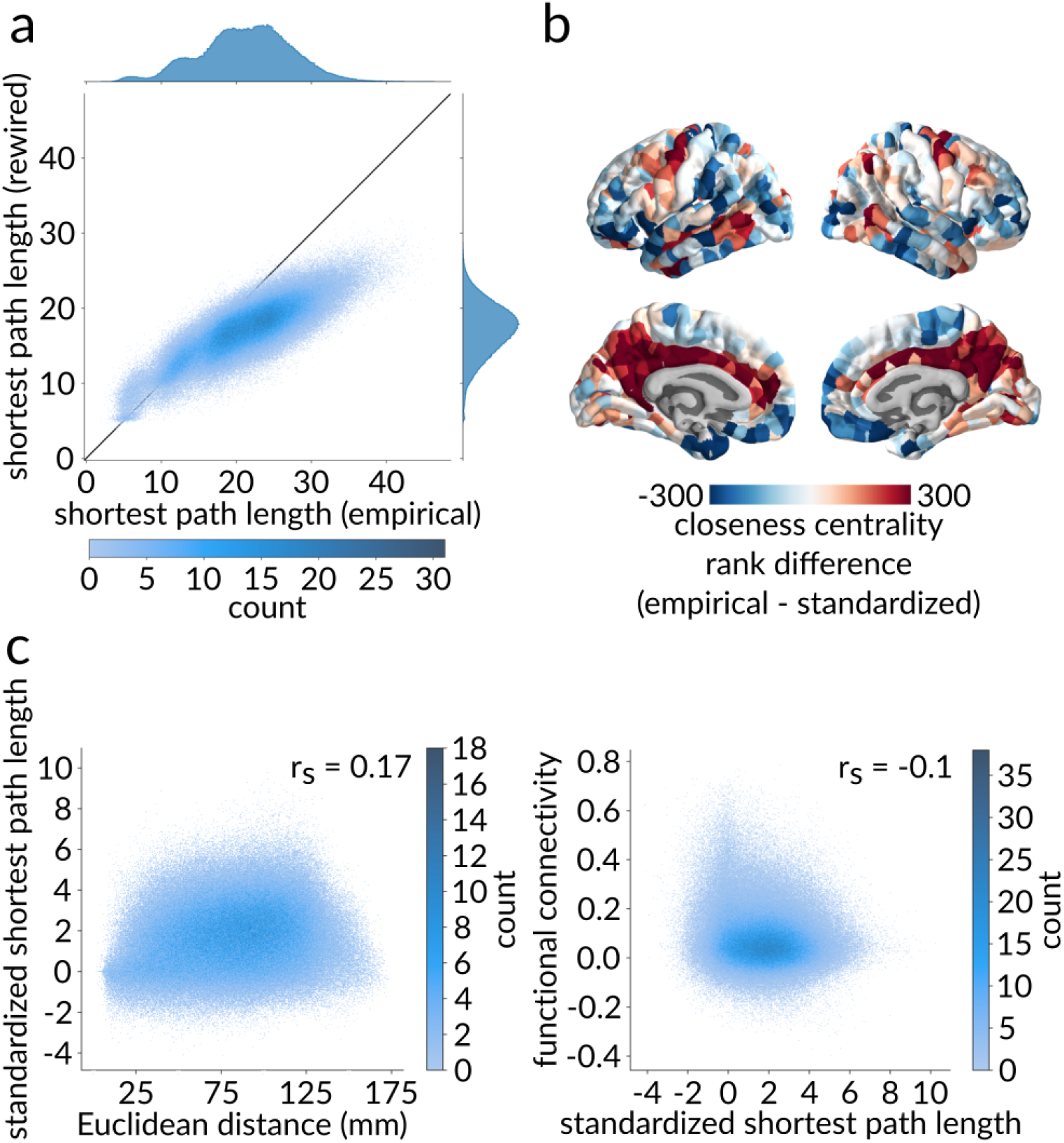
Disentangling the contributions of topology and geometry. (a) Scatter plot between empirical (abscissa) and rewired (ordinate) shortest path lengths obtained using a geometry-preserving null model, where each point represents a pair of brain regions. Marginal distribution histograms are shown on the top and right axes. Points that appear below the identity line correspond to paths with a shorter length in the rewired networks than in the empirical network, and vice versa for points above the identity line. (b) Brain map of the region-wise differences between rank-transformed closeness centrality (inverse mean path length to the rest of the network) computed using empirical and standardized shortest path lengths. Red regions are more integrated in the empirical network, and blue regions are more integrated in the standardized network. (c) Relationships between standardized shortest path length and Euclidean distance (left) and functional connectivity (right). As expected due to the edge length-preserving surrogate model, the relationship between standardized shortest path length and Euclidean distance is considerably attenuated compared to the result obtained using strictly topology-preserving nulls. The relationship between standardized shortest path length and functional connectivity is maintained (*r*_s_ = −.1, *p* ≈ 0), but is not exponential anymore.

**Figure S3.**
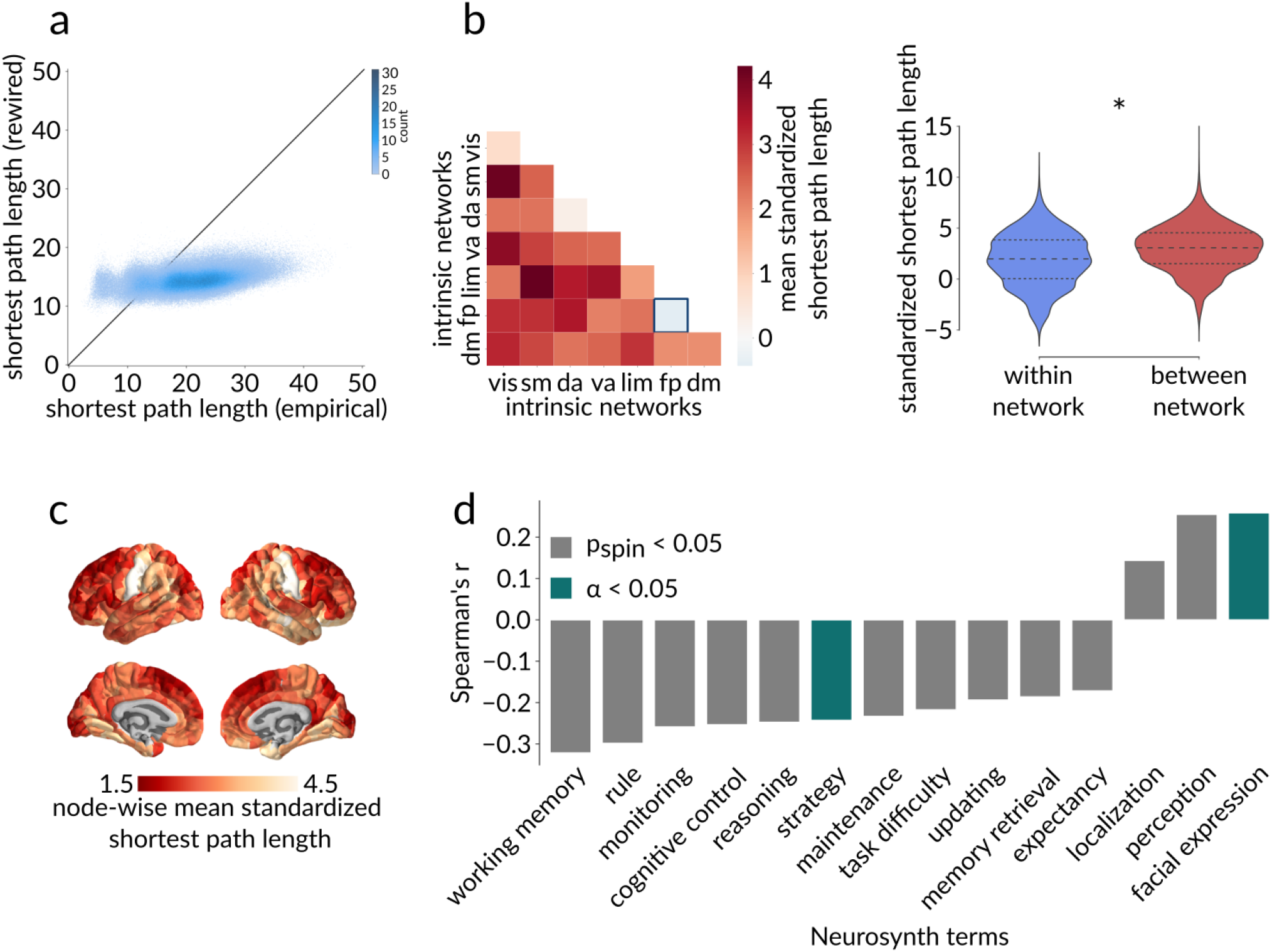
Sensitivity analysis - *Validation* dataset. (a) Scatter plot between empirical (abscissa) and rewired (ordinate) shortest path lengths obtained in the *Validation* sample, where each point represents a pair of brain regions. (b) Left: Heatmap of the mean standardized path lengths across node pairs belonging to the same intrinsic network (diagonal) and to different intrinsic networks (off-diagonal). The frontoparietal network’s greater-than-expected internal communication capacity is replicated in the *Validation* dataset. Right: The mean within-network standardized path length is also significantly shorter than the mean between-network standardized path length in the *Validation* dataset (*p*_spin_ < .001). (c) Brain map of mean standardized path length from each node to the rest of the network from the *Validation* set, with red denoting a greater integration of the node within the network and yellow denoting a lower integration. (d) Significant anticorrelations (*p*_spin_ < .05 in grey; Bonferroni corrected, **α = .05** in green) between node-wise mean standardized path length and Neurosynth functional activation maps associated to higher-order cognitive functions were replicated in the *Validation* dataset.

**Figure S4.**
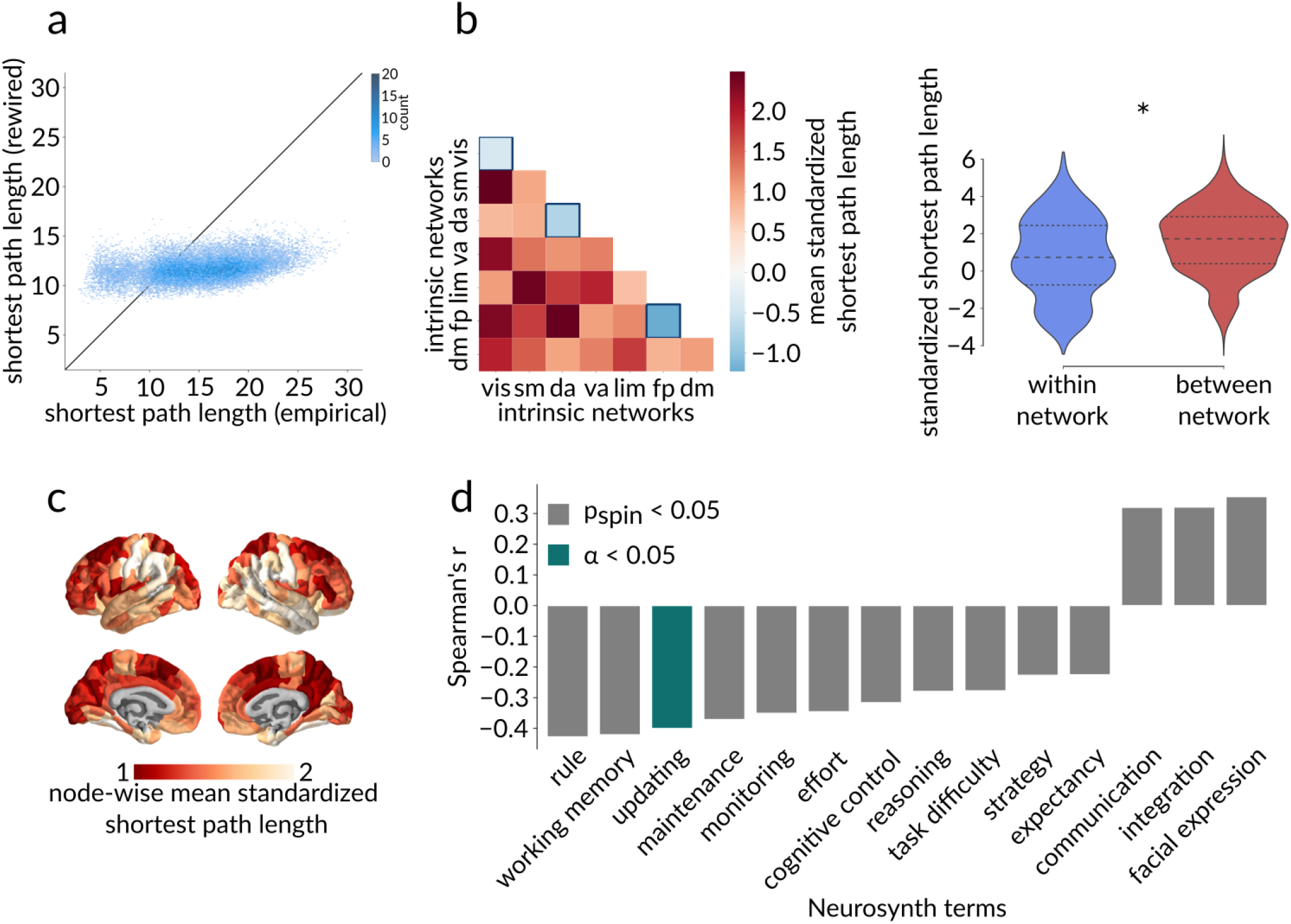
Sensitivity analysis - 219 nodes resolution. (a) Scatter plot between empirical (abscissa) and rewired (ordinate) shortest path lengths obtained using the lower resolution Cammoun parcellation, where each point represents a pair of brain regions. (b) Left: Heatmap of the mean standardized path lengths across node pairs belonging to the same intrinsic network (diagonal) and to different intrinsic networks (off-diagonal). In addition to replicating the frontoparietal network’s greater-than-expected internal communication capacity, this partition also identifies the communication pathways internal to the visual and the dorsal attention networks as displaying greater-than-expected efficiencies. Right: The mean within-network standardized path length is also significantly shorter than the mean between-network standardized path length when using the 219 nodes resolution of the Cammoun atlas (*p*_spin_ < .001). (c) Lower resolution brain map of mean standardized path length from each node to the rest of the network, with red denoting a greater integration of the node within the network and yellow denoting a lower integration. (d) Significant anticorrelations (*p*_spin_ < .05 in grey; Bonferroni corrected, **α = .05** in green) between node-wise mean standardized path length and Neurosynth functional activation maps associated to higher-order cognitive functions were replicated at a lower resolution.

**Figure S5.**
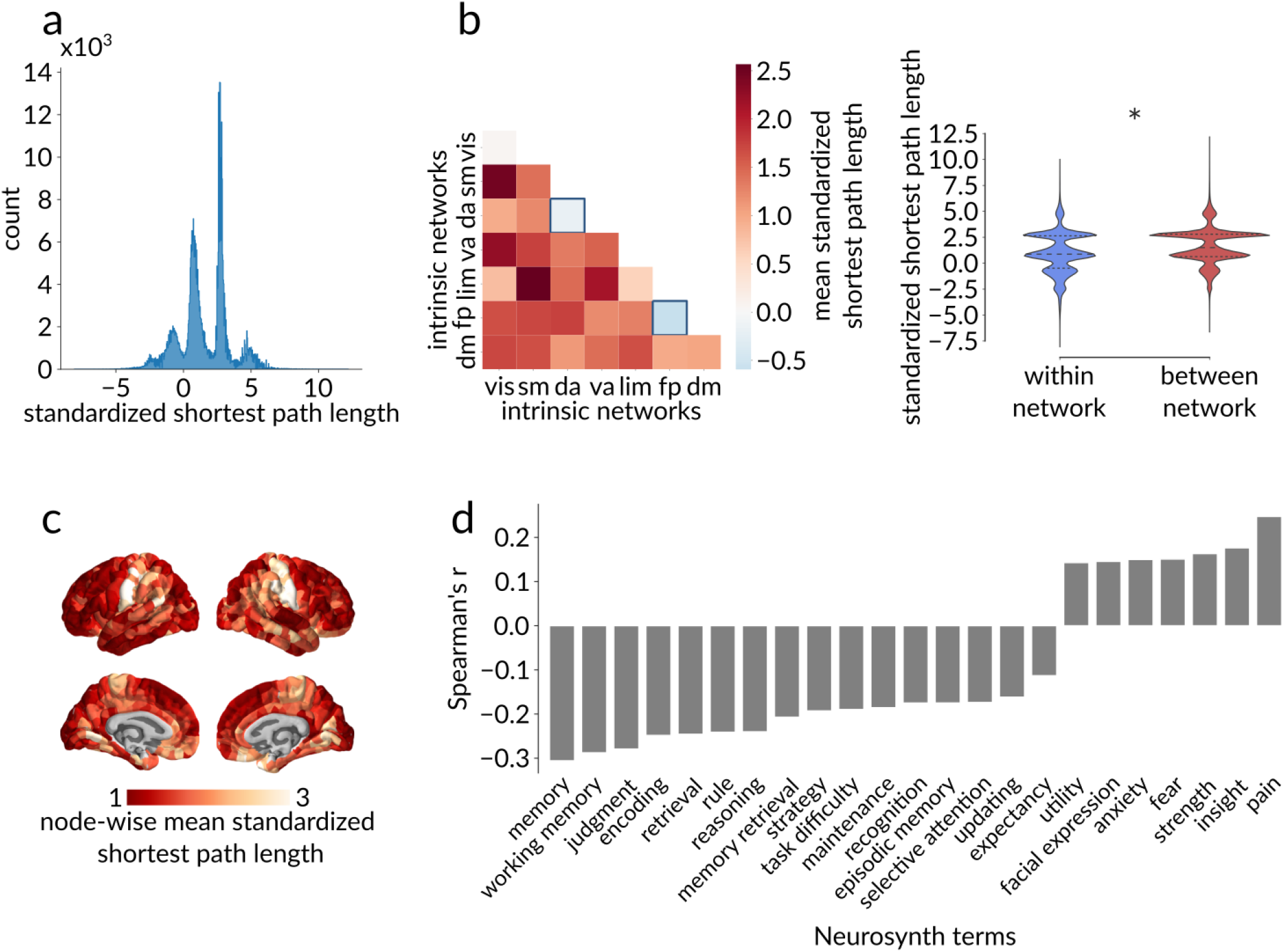
Sensitivity analysis - binary structural networks. (a) Distribution of standardized binary shortest path lengths (z-scores) computed from a binary group-consensus structural network for all pairs of brain regions. (b) Left: Heatmap of the mean standardized path lengths across node pairs belonging to the same intrinsic network (diagonal) and to different intrinsic networks (off-diagonal). In addition to replicating the frontoparietal network’s greater-than-expected internal communication capacity, binary path lengths also identify greater-than-expected communication capacity in the dorsal attention network. Right: The mean within-network standardized path length is also significantly shorter than the mean between-network standardized path length when using binary structural networks (*p*_spin_ < .001). (c) Brain map of mean standardized binary path length from each node to the rest of the network, with red denoting a greater integration of the node within the network and yellow denoting a lower integration. (d) Significant anticorrelations (*p*_spin_ < .05) between node-wise mean standardized path length and Neurosynth functional activation maps associated to higher-order cognitive functions were replicated using binary structural networks.

**Figure S6.**
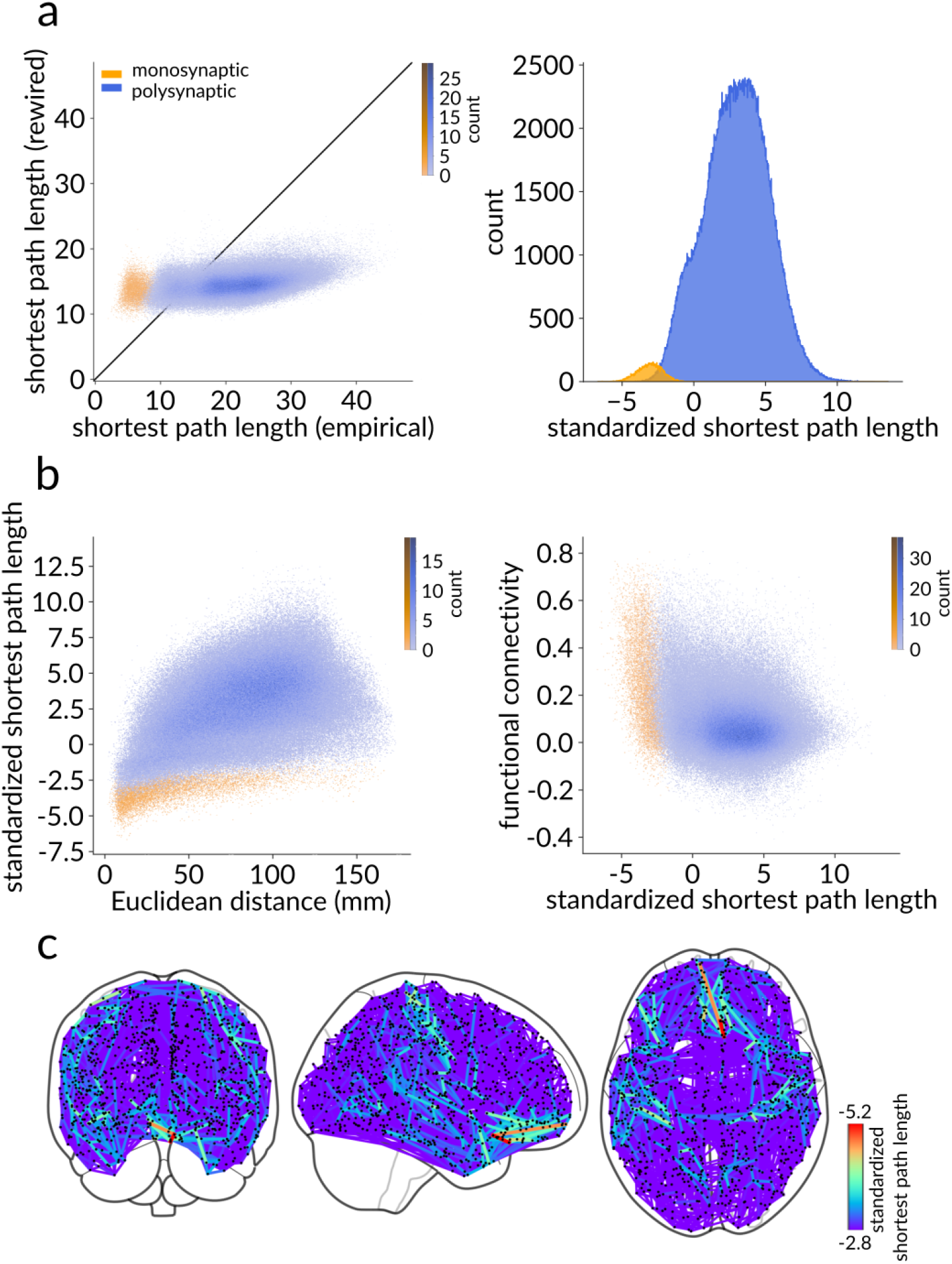
Sensitivity analysis - Polysynaptic communication pathways. (a) Left: Scatter plot between empirical (abscissa) and rewired (ordinate) shortest path lengths. Each point represents a pair of brain regions. Right: Distribution of standardized shortest path lengths (z-scores) for all pairs of brain regions. As expected, a large proportion of negative standardized path lengths are attributed to monosynaptic pathways. Monosynaptic communication pathways appear in yellow, whereas polysynaptic shortest paths are coloured in blue. (b) Relationships between standardized shortest path length and Euclidean distance (left) and functional connectivity (right), with monosynaptic connections identified in yellow and polysynaptic paths in blue. (c) Spatial distribution of the top 1% unexpectedly short polysynaptic path lengths

**Figure S7.**
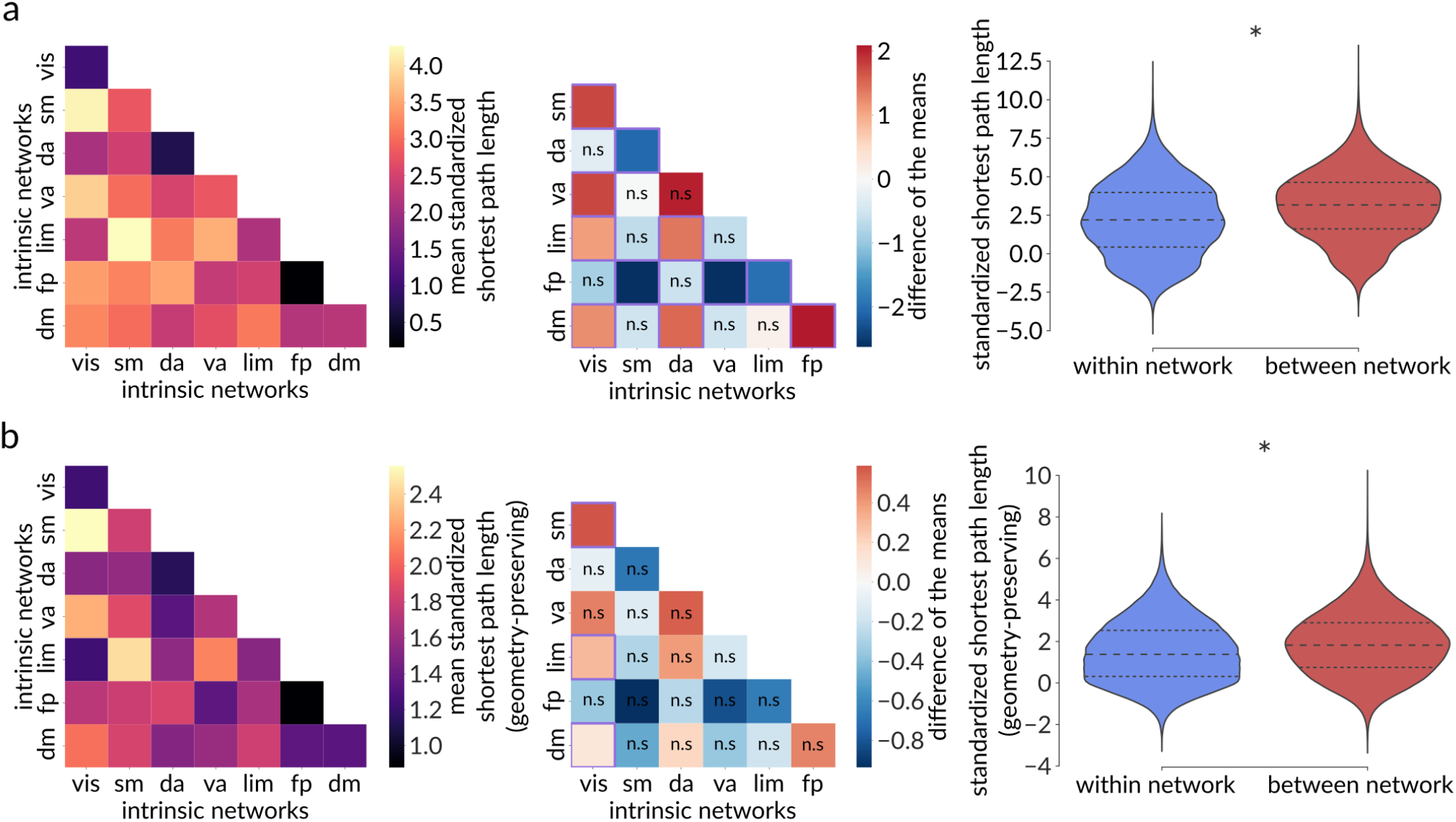
Sensitivity analysis - Polysynaptic communication pathways delineate functional systems. (a) Left: Heatmap of the mean standardized path lengths across node pairs belonging to the same intrinsic network (diagonal) and to different intrinsic networks (off-diagonal). In comparison to the previous results considering all communication pathways, including monosynaptic connections, the frontoparietal network’s mean standardized path length is no longer negative, suggesting that its greater-than expected internal communication capacity is partly due to highly efficient direct anatomical connections. Middle: Heatmap of the pairwise differences of the means among Yeo intrinsic networks, calculated as the mean value of the network on the y-axis minus the mean value of the network on the x-axis, with the mean value corresponding to the mean standardized path length across node pairs belonging to the same network (diagonal elements of the left heatmap). A purple square indicates significant difference of the means based on network label permutation using spatial autocorrelation-preserving null models (Bonferroni corrected, ***α*** = **.05**), whereas “n.s.” denotes not significant differences. This plot recapitulates the results obtained when taking all communication pathways into account, with the frontoparietal network displaying the highest internal communication capacity and the somatomotor network exhibiting the lowest internal communication capacity. Right: The mean within-network standardized path length is also significantly shorter than the mean between-network standardized path length when considering only polysynaptic communication pathways (*p*_spin_ < .001). (b) Same as (a) but for polysynaptic communication pathways standardized using a geometry-preserving null model.

**Figure S8.**
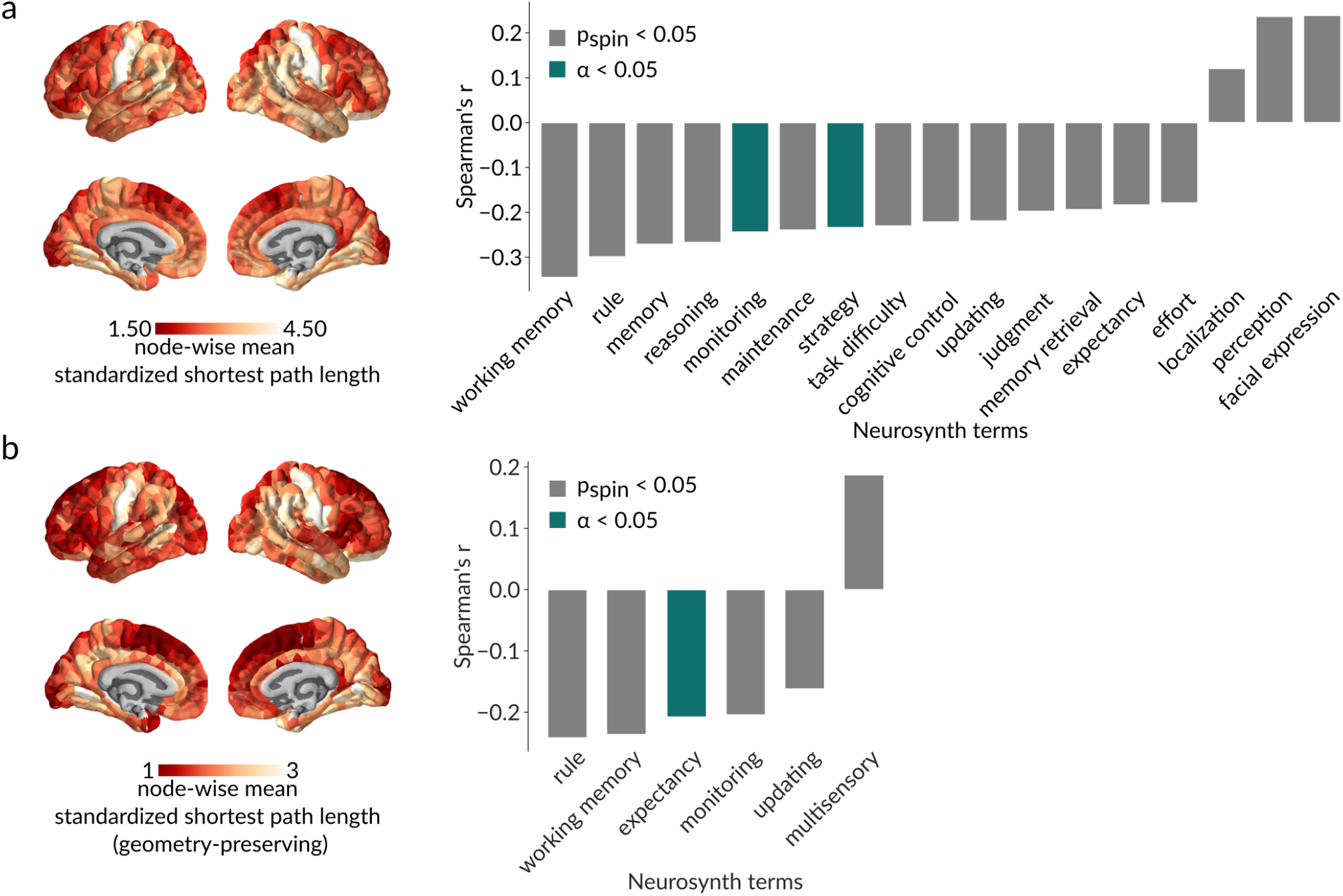
Sensitivity analysis - Polysynaptic communication capacity and functional specialization. (a) Left: Brain map of mean standardized polysynaptic path length from each node to the rest of the network, with red denoting a greater integration of the node within the network and yellow denoting a lower integration. Right: Significant anticorrelations (*p*_spin_ < .05 in grey; Bonferroni corrected, **α = .05** in green) between node-wise mean standardized path length and Neurosynth functional activation maps associated to higher-order cognitive functions are replicated in polysynaptic communication pathways. (b) Same as (a) but for polysynaptic communication pathways standardized using a geometry-preserving null model.

**Figure S9.**
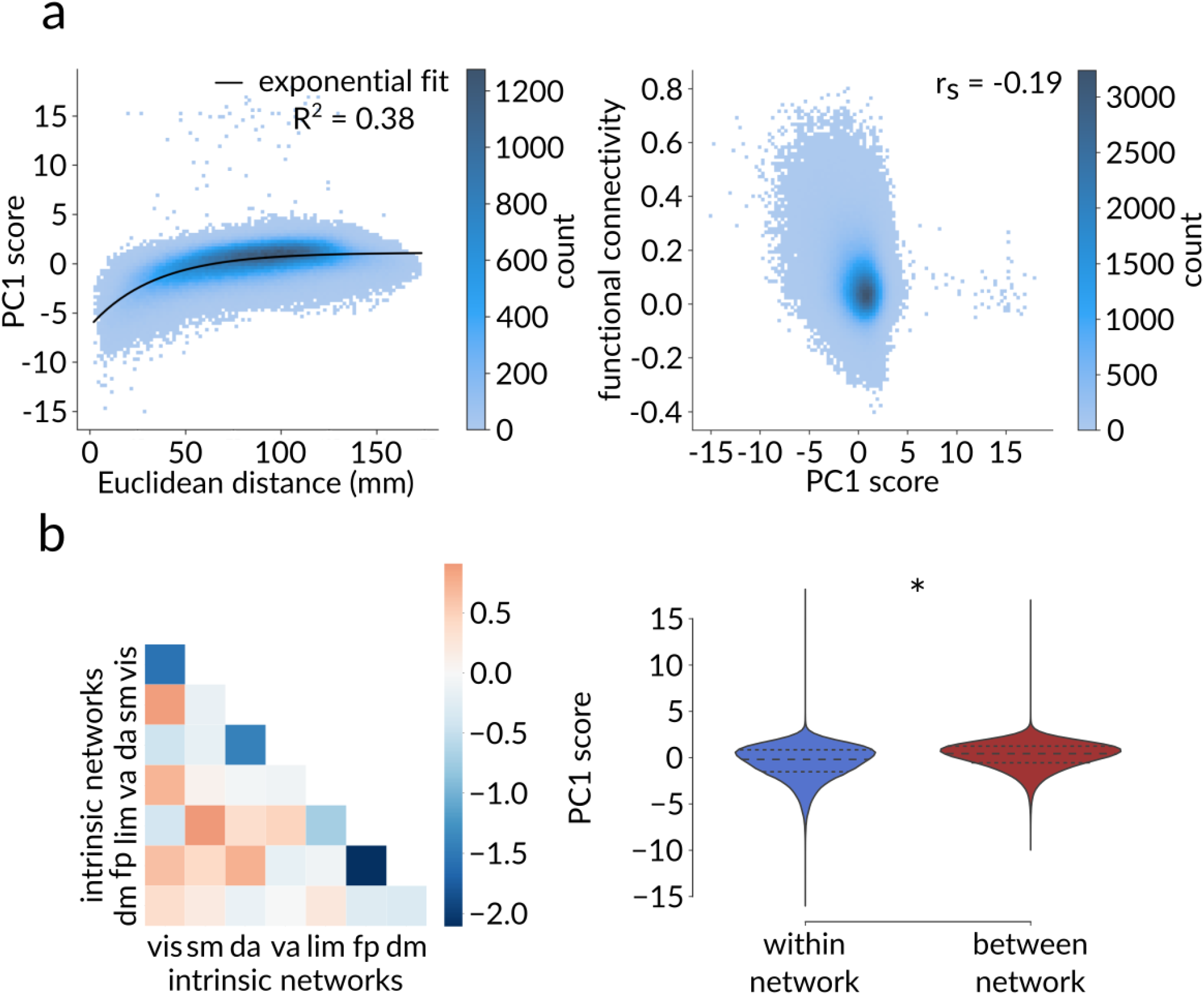
Sensitivity analysis - Polysynaptic communication pathways from multiple communication models. The PC1 aggregate communication score recapitulates the results obtained using standardized shortest path length even if we circumscribe the analyses to PC1 scores of node pairs separated by more than one synapse. (a) Left: Growth of the PC1 score as a function of Euclidean distance. The black line corresponds to the fitted exponential ***y*** = −**7.49e**^-0.03*x*^ + **1.12**. Right: Negative Spearman correlation between functional connectivity and PC1 score (*r*_s_ = −.19, *p* ≈ 0). (b) Left: Heatmap of the mean PC1 score across node pairs belonging to the same intrinsic network (diagonal) and to different intrinsic networks (off-diagonal). Right: Significantly lower within-network than between-network PC1 score (*p*_spin_ < .001).

**Figure S10.**
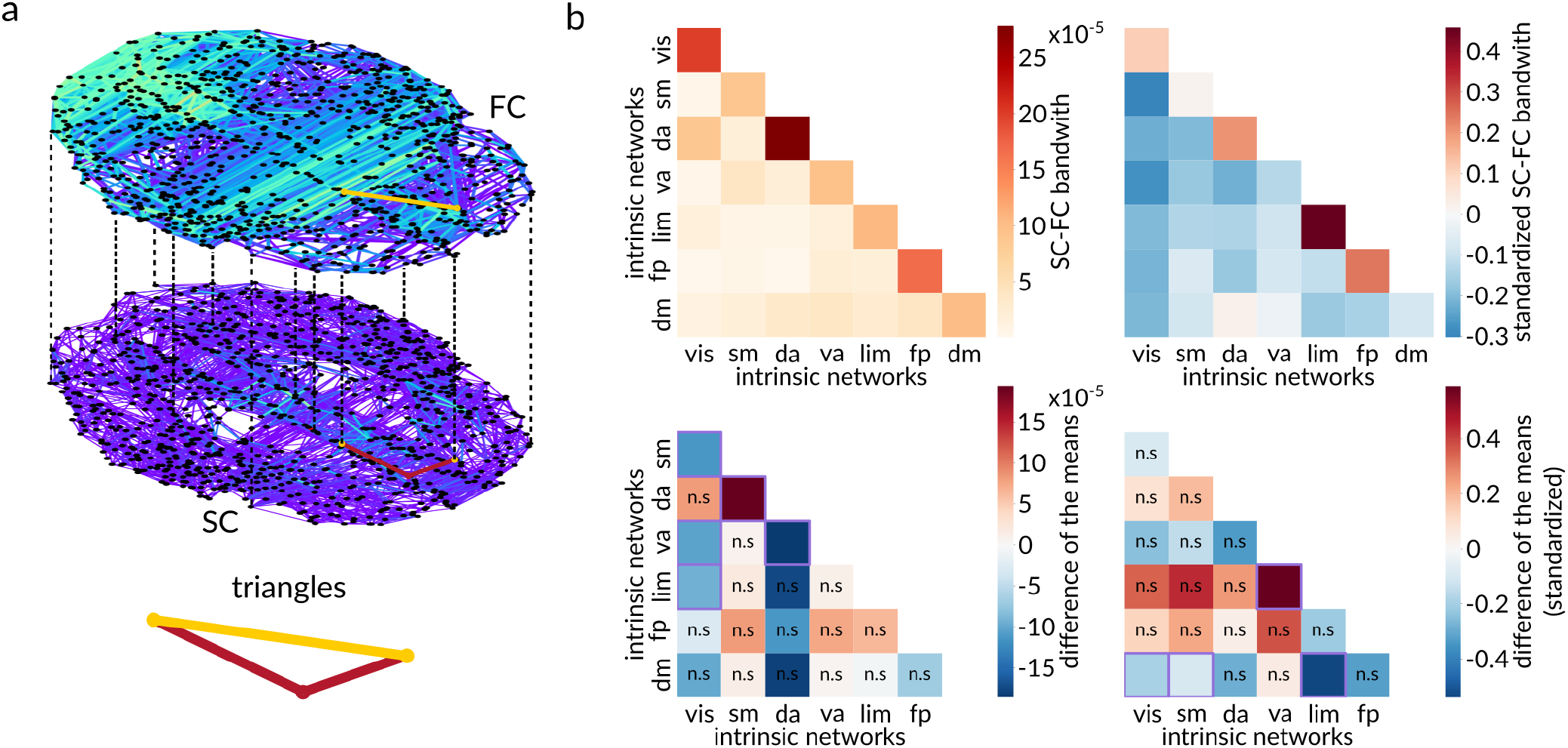
Sensitivity analysis - higher-order connectivity. (a) 2-layer multiplex of cortical connectivity. Edges are showed in red in the lower structural connectivity (SC) layer, whereas they appear in yellow in the upper functional connectivity (FC) layer. Dashed lines represent pseudo-edges connecting the two layers to close structural pathways by an FC edge, forming SC-FC polygons. (b) Top: Heatmap of the mean empirical (left) and standardized (right) SC-FC triangles bandwidth across node pairs of the FC layer belonging to the same intrinsic network (diagonal) and to different intrinsic networks (off-diagonal). Bottom: Heatmap of the pairwise differences of the means among Yeo intrinsic networks, calculated as the mean value of the network on the y-axis minus the mean value of the network on the x-axis, with the mean value corresponding to the mean empirical (left) and standardized (right) SC-FC triangles bandwidth across node pairs of the FC layer belonging to the same network (diagonal elements of the top heatmap). A purple square indicates significant difference of the means based on network label permutation using spatial autocorrelation-preserving null models (Bonferroni corrected, ***α*** = **.05**), whereas “n.s.” denotes not significant differences. The dorsal attention network displays a consistently higher internal SC-FC triangles bandwidth compared to other networks, whereas the somatomotor network exhibits a consistently lower SC-FC triangles bandwidth. These results are not maintained when standardizing SC-FC bandwidth, with the limbic network now showing the highest triangles bandwidth, and the ventral attention network showing the lowest.

## Notes

https://github.com/fmilisav/milisav_dyadic_communication

### Competing Interest Statement

The authors have declared no competing interest.

### Summary of Updates

Introduction extended; Parts of the Supplementary Materials incorporated into the main body of the manuscript; New sensitivity analyses added.

